# Collagen VI is a conserved pro-fibrotic signal disrupting muscle regeneration across distinct human myopathies

**DOI:** 10.1101/2025.10.17.683021

**Authors:** Laura Muraine, Mona Bensalah, Stephen Gargan, Paul Dowling, Anne Bigot, Valérie Allamand, Jamila Dhiab, Maria Kondili, Sophie Perié, Jean Lacau St-Guily, Gillian Butler-Browne, Vincent Mouly, Kay Ohlendieck, Capucine Trollet, Elisa Negroni

## Abstract

Muscle fibrosis is a major driver of progression in diverse myopathies, yet the conserved molecular mediators of this process in humans remains poorly defined. Here, we identify collagen VI as a common pro-fibrotic and regeneration-impairing extracellular matrix (ECM) component across three distinct human myopathies: Duchenne Muscular Dystrophy (DMD), Oculopharyngeal Muscular Dystrophy (OPMD), and Inclusion Body Myositis (IBM). Proteomic profiling of fibrotic biopsies revealed consistent upregulation of collagen VI and laminin γ1, alongside disease-specific alterations involving other ECM proteins. Fibro-adipogenic progenitors (FAPs) are the predominant source of these ECM components, including collagen VI and laminin γ1. Functionally, xenotransplantation of patient-derived FAPs into regenerating mouse muscle induced localized collagen deposition, myofiber atrophy, and depletion of Pax7⁺ muscle stem cells. Mechanistic assays demonstrated that FAP-derived collagen VI is sufficient to impair myogenic fusion, while silencing COL6 in patient FAPs restores fusion capacity, directly linking pathological collagen VI deposition to regeneration failure. Our findings uncover collagen VI as a conserved effector of fibrosis and stem cell niche disruption in human myopathies, positioning it as a potential therapeutic target across genetically and clinically distinct muscle diseases.

## Introduction

Muscular disorders encompass a diverse group of inherited and acquired diseases characterized by progressive muscle weakness and degeneration. These conditions differ in age of onset, affected muscle groups and severity ^1^. Duchenne muscular dystrophy (DMD), a severe genetic myopathy caused by primary defects in the X-chromosomal *DMD* gene, leads to widespread muscle degeneration from early childhood ^2^. In contrast, oculopharyngeal muscular dystrophy (OPMD) and inclusion body myositis (IBM) are late-onset myopathies typically manifesting after the age of 50, with more localized muscle involvement. A common feature of OPMD and IBM is pharyngeal muscle weakness, leading to dysphagia ^3,4^. OPMD is caused by a triplet repeat expansion mutation, whereas IBM is an acquired inflammatory myopathy.

A hallmark of these myopathies is muscle fibrosis, characterized by an excessive deposition of extracellular matrix (ECM) components ^5^. Over time, ECM replaces muscle fibers, compromising tissue integrity and limiting the efficacy of therapeutic interventions. Recent single-cell omics studies have provided insights into the cellular heterogeneity underlying dystrophic progression ^6,7^. While intercellular communication is recognized as a key factor in fibrosis, comprehensive analyses of ECM composition in human myopathies remain limited.

The ECM is a dynamic scaffold that regulates tissue development, structural integrity, and key cellular functions, such as proliferation, survival, and stem cell maintenance ^8–11^. Remodeling of the matrisome, which encompasses both structural ECM components (core matrisome) and regulatory proteins ^12^, plays a central role in the apparition and maintenance of fibrosis. The key ECM-producing cells in skeletal muscle are fibroadipogenic progenitors (FAPs). FAPs are involved in both physiological and pathological processes. However, their role in human skeletal muscle fibrosis remains poorly defined. Understanding quantitative and qualitative changes in the matrisome and the exact contribution of FAPs to the interstitial matrix composition is essential for deciphering the mechanisms driving skeletal muscle fibrosis and developing corrective strategies.

In this study, we performed a comparative proteomic study by liquid chromatography coupled to mass spectrometry (LC-MS/MS) of three distinct human myopathies involving fibrosis: DMD, OPMD, and IBM. By comparing affected fibrotic muscles from patients to those from healthy age-matched subjects, we identified matrisome alterations in each pathology. We defined different ECM remodeling signatures across myopathies but a shared over-abundance of collagen VI. Furthermore, we isolated FAPs from patient biopsies and defined their secretion of key ECM proteins *in vitro* and *in vivo*. Beyond ECM remodeling, our results highlight the impact of FAP-driven ECM alterations on muscle regeneration. Indeed, *in vivo* experiments demonstrated that FAP-secreted ECM components, including Collagen VI and basal lamina proteins, influence muscle stem cell behavior and muscle tissue regenerative capacity. Silencing COL6 in patient FAPs was able to restore fusion capacity, directly linking pathological collagen VI deposition to regeneration failure. This work advances our understanding of ECM regulation in skeletal muscle diseases and uncover collagen VI as a conserved effector of fibrosis and a potential therapeutic target across genetically and clinically distinct muscle diseases.

## Results

### Fibrotic FAPs share similar features across DMD, OPMD and IBM

Muscle fibrosis is a hallmark of many myopathies, yet it remains unclear whether its cellular and molecular drivers are conserved across diseases or if each myopathy presents a distinct fibrotic signature. To investigate this, we analyzed three myopathies characterized by the presence of fibrosis: Duchenne Muscular Dystrophy (DMD), Oculopharyngeal muscular Dystrophy (OPMD) and Inclusion Body Myositis (IBM). Biopsies from affected muscles exhibited severe fibrosis covering 38% to 69% of the total muscle section, compared to a median of 9% in healthy muscles (CTL) **(Fig. 1A).** They also showed 13% to 23% of regenerating fibers positive for embryonic myosin heavy chain (MyHC-emb) **(Fig. 1B)**, along with a high abundance of fibroadipogenic progenitors (FAPs), identified as interstitial cells expressing the CD90 marker **(Fig. 1C).** To characterize the cellular actors of fibrosis and assess potential disease-specific differences, we isolated FAPs from fibrotic muscles of all three myopathies and analyzed their properties *in vitro*. FAPs isolated from affected muscles all displayed increased proliferation rates compared to those from non-fibrotic control muscles **(Fig. 1D)**. Moreover, when co-cultured with myotubes **(Fig. 1E)**, disease-derived FAPs impaired myogenic fusion, resulting in a reduced fusion index, an effect not observed in the presence of FAPs from control muscles **(Fig. 1F-G)**.

**Fig. 1.**
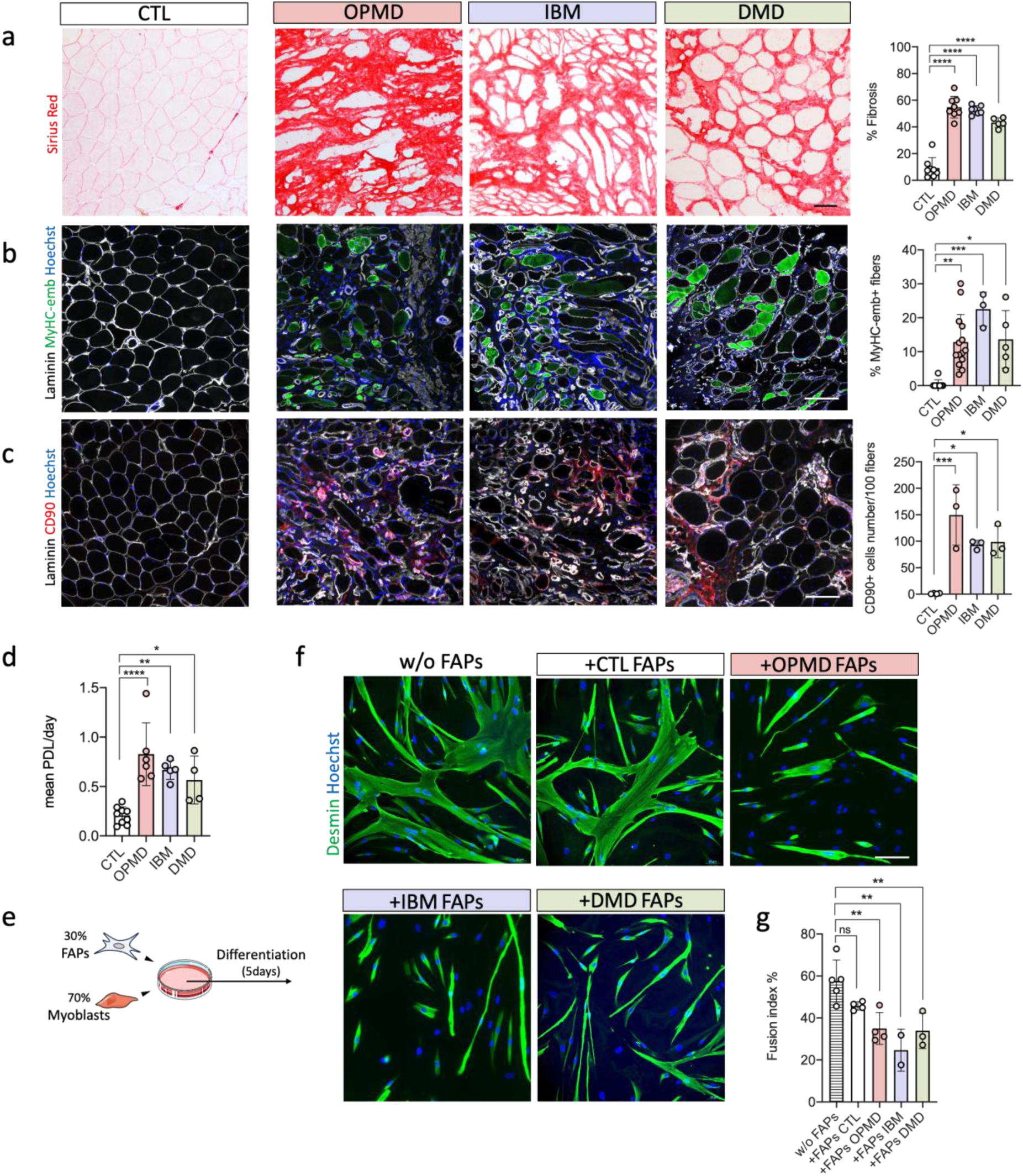
FAPs are abundant in affected muscles of OPMD, IBM and DMD patients and impair myotubes fusion *in vitro*. **(a)** Sirius red coloration of human skeletal muscle biopsies from healthy CTL, OPMD, IBM and DMD patients. Scale bar: 50μm - Quantification of the percentage of fibrosis by Sirius red coloration (n=5-8 biopsies per condition). **(b)** Immunostaining of laminin (white), Hoechst (blue) and embryonic myosin heavy chain (MyHC-emb (MYH3)) (green). Scale bar: 50μm - Quantification of the percentage of MyHC-emb positive fibers for each condition (n=3-14 biopsies per condition). **(c)** Immunostaining of laminin (white), Hoechst (blue) and CD90 (red) on human biopsies - Quantification of the number of interstitial cells positive for CD90 per 100 fibers (n=3-4 biopsies per condition). **(d)** *In vitro* proliferation rate expressed as the mean population doubling (PDL) /day of FAPs isolated from healthy CTL, OPMD, IBM and DMD muscle biopsies. **(e)** Schematic representation of coculture protocol; FAPs are co-cultured with myoblasts at a 30%/70% ratio for 5 days in differentiation medium. **(f)** Immunostaining of myotubes/FAPs co-cultures (Desmin, green; Hoechst, blue) after 5 days of differentiation, scale bar: 50μm - Quantification of the fusion index of the myoblasts, with or without FAPs. Data are presented as mean ±SD, with P-values obtained by ordinary one-way ANOVA test followed by Tukey’s multiple comparisons test (*P < 0.1 **P < 0.01, ***P < 0.001, ****P < 0.0001).

### Mass spectrometry analysis reveals disease-specific and shared changes in muscle proteome

To investigate fibrosis-associated ECM remodeling in DMD, OPMD and IBM, we analyzed the proteome of fibrotic muscle biopsies of each pathology by LC-MS/MS. Each patient’s muscles were compared to age-matched control muscles. In total, 172 proteins exhibited a significantly different abundance compared to control, with 32 proteins increased in OPMD, 45 in IBM and 133 (77 increased, 56 decreased) in DMD samples **(Fig. 2A and Tables 1-3)**, suggesting more pronounced proteomic alterations in DMD when compared to age-matched healthy muscles. Of these, 25 proteins were shared between two of the three myopathies, while only six proteins were commonly differentially expressed across all three diseases **(Fig. 2A and supplementary Table 1),** highlighting both shared and disease-specific signature. To characterize disease-specific proteomes, we performed gene ontology biological process enrichment analyses. Among the most enriched terms, several were related to ECM organization, highlighting the expected increase in ECM protein abundance in pathological biopsies. Additional pathway-level analysis revealed molecular features reflecting the distinct pathophysiology of each disorder. OPMD and IBM biopsies were primarily enriched in pathways linked to mitochondrial dysfunction and protein folding stress responses, including upregulation of molecular chaperones and oxidative phosphorylation components, suggesting cellular stress and impaired proteostasis. DMD biopsies, by contrast, were distinguished by alterations in cytoskeletal organization, chronic inflammatory signaling, and glycolytic metabolism, pointing to ongoing regeneration and immune activation **(Fig. 2B)**. Using Outcyte prediction tool ^13^, we determined that among differentially expressed proteins, more than 68% for OPMD, 46% for IBM and 60% for DMD were annotated as secreted, conventionally or unconventionally **(Fig. 2C and Tables 1-3).**

**Fig. 2.**
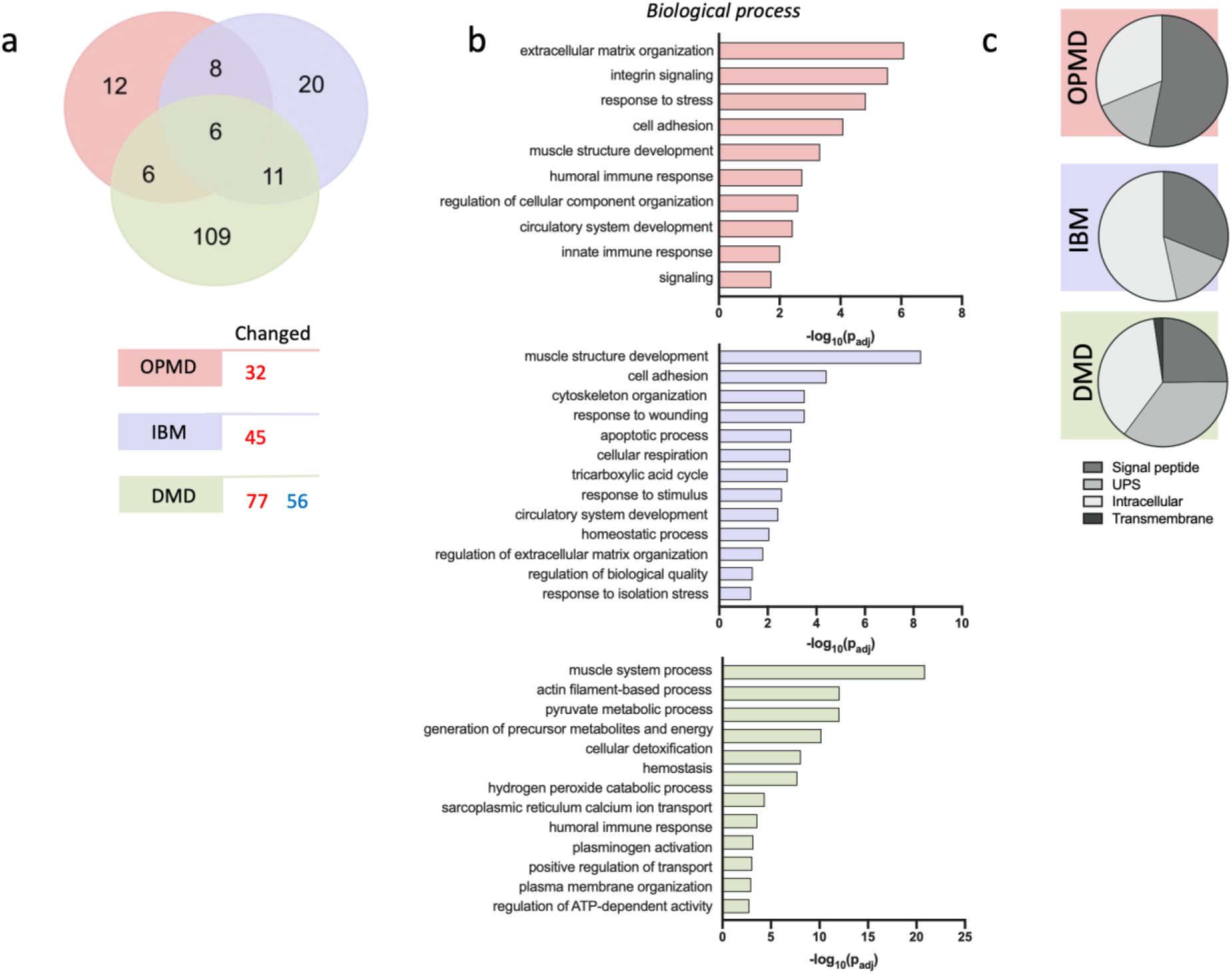
Proteomic analysis highlights different disease profiles and a shared ECM signature. **(a)** Number of differentially expressed proteins identified among pathologies when compared to their age-matched control muscles (increased in red, decreased in blue)- Venn diagram representing differentially expressed proteins shared between myopathies. **(b)** Significantly altered proteins were subjected to a GO term analysis to identify associated biological processes using gProfiler. Bar charts of top enriched terms for each comparison. **(c)** Representation of the repartition of proteins on their predicted secretion potential using Outcyte prediction tool (UPS; unconventional protein secretion).

### FAPs drive core matrisome remodeling across myopathies

To further dissect fibrosis-associated ECM remodeling, we filtered the identified proteins using the human matrisome database ^14^, which categorizes ECM components into the “core matrisome” (collagens, glycoproteins and proteoglycans) and “matrisome associated” proteins (ECM regulators, ECM-affiliated proteins and secreted factors). Between 18% and 43 % of the differentially expressed proteins were ECM-related, with the majority belonging to the core matrisome **(Fig. 3A, 3B and table 4A-D)**. We identified five core matrisome proteins consistently altered across all three myopathies: Collagen VI (COL6A1, COL6A2 and COL6A3), laminin subunit gamma 1 (LAMC1) and perlecan (HSPG2) **(Fig. 3A, 3C)**. Notably, HSPG2 levels diverged showing decreased expression in DMD but increased expression in OPMD and IBM **(Fig. 3C)**. Disease-specific ECM alterations included upregulation of collagen alpha-1(I) chain (COLIA1) and fibronectin 1 (FN1), the latter being significantly upregulated in both OPMD and IBM and exhibiting the highest fold change **(Fig. 3C and table 4A-B).** ECM remodeling proteins were also dysregulated, with alpha-2-macroglobulin (A2M) and alpha-1-antitrypsin (SERPINA1) increased in DMD, antithrombin-III (SERPINC1) altered in both DMD and OPMD or plasminogen activation system (PLG) altered in both OPMD and IBM. Shared dysregulation of ECM degradation pathways suggests a common impairment in ECM turnover across diseases. Changes in annexins, which are involved in membrane repair, included ANXA1 increased in IBM, ANXA2 in DMD and OPMD, and ANXA5 and ANXA6 in DMD and IBM. Basal membrane proteins such as COL4A1 and laminin subunit beta-2 (LAMB2) were specifically upregulated in OPMD **(Fig. 3A, 3C).** Immunofluorescence analysis **(Fig. 3D)** further validated the presence of high level of Collagen VI across all three myopathies. Altogether, these findings reveal a combination of shared ECM remodeling features and distinct disease-specific signatures across myopathies. To further delineate the cellular sources of ECM proteins, we analyzed a publicly available human single-cell RNA sequencing (scRNA-seq) dataset from ^15^. This analysis confirmed that FAPs are the predominant source of the identified ECM proteins, including collagens, glycoproteins and basal lamina proteins **(Fig. 3E)**. To investigate their role in ECM production, FAPs isolated from control, DMD, OPMD, and IBM muscle samples were cultured *in vitro* and found to actively secrete key ECM components, forming networks of collagen VI fibrils, glycoproteins (Tenascin X, Fibronectin) and basal lamina components (Collagen IV) **(Fig. 3F)**.

**Fig. 3.**
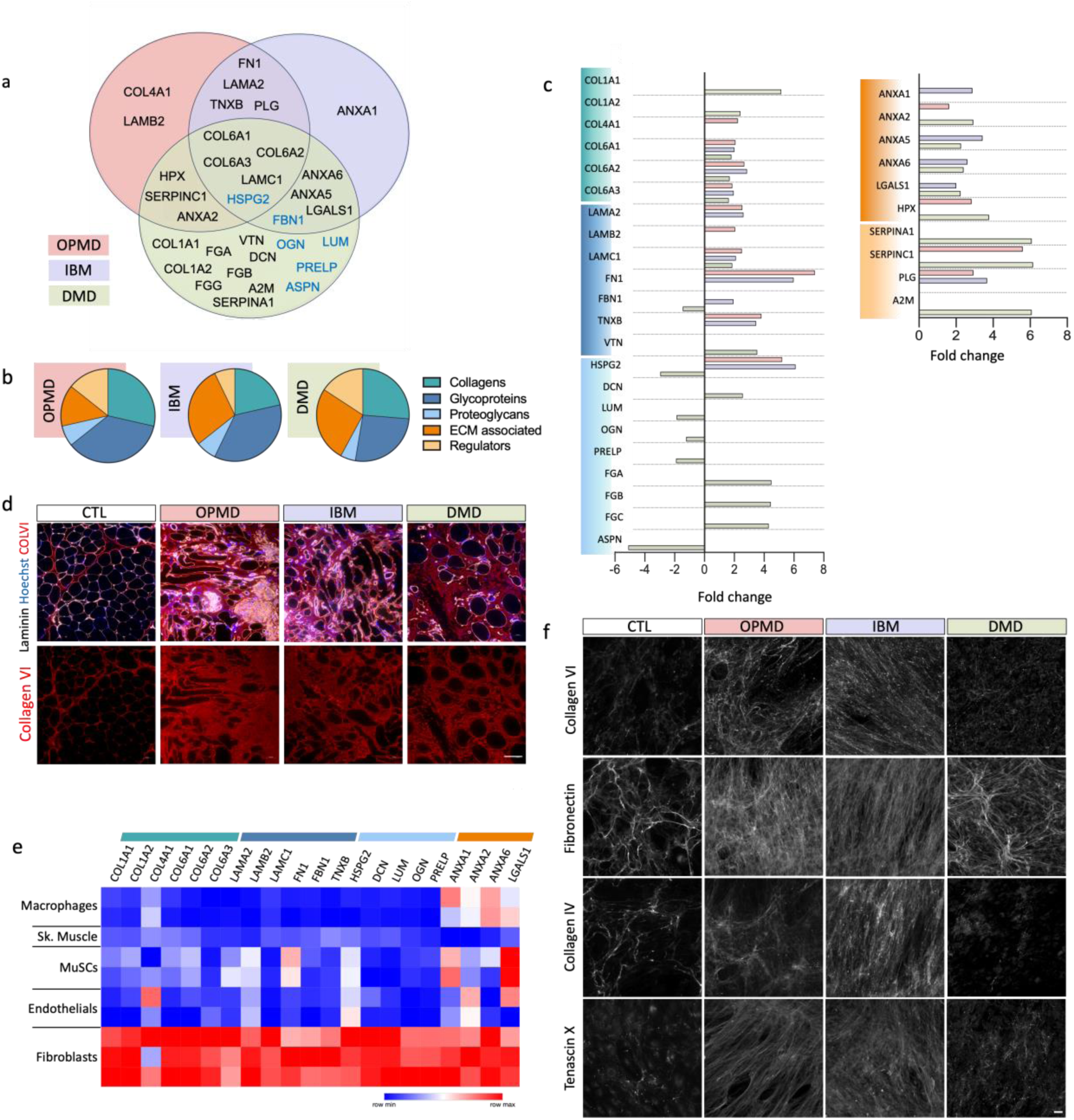
FAPs are the main actors responsible for matrisome alterations in myopathies. **(a)** Venn diagram representing matrisome proteins (from the matrisome database matrisomeproject.mit.edu) differentially expressed between pathologies and their age-matched control muscles. In blue, proteins decreased in DMD biopsies. (**b)** Representation of the repartition of the differentially expressed proteins for each group of ECM related proteins, in each pathology. **(c)** Representation of the fold changes in abundance of differentially expressed ECM-associated proteins for each pathology (same color code as 3A) compared to their controls. Color code for proteins corresponds to the matrisome database protein groups shown in Fig 3B. **(d)** Immunostaining of laminin (white), Hoechst (blue) and Collagen alpha-3(VI) chain (red). Scale bar: 50μm **(e)** Heatmap of the expression of ECM proteins in several resident cell types within skeletal muscle on sc-RNA-seq dataset extracted from De Micheli *et al* 2020 (GSE143704). MuSCs: Muscle stem cells; Sk muscle: skeletal muscle **(f)** Immunostaining of Collagen alpha-3(VI) chain, fibronectin, Collagen IV and Tenascin X on ECM produced by FAPs *in vitro*. Scale bar: 50μm

### FAPs-driven ECM modifications impact muscle regeneration and stem cell niche maintenance

To assess the functional impact of disease-derived FAPs on muscle regeneration, we injected human FAPs isolated from fibrotic muscle biopsies into the regenerating *tibialis anterior* (TA) muscle of Rag2^-/-^ Il2rb^-/-^ immunodeficient mice **(Fig. 4A)**. Four weeks post-injection, following complete regeneration, we detected human FAPs in the interstitial space between newly formed muscle fibers using human-specific antibodies **(Fig. 4B)**. All FAPs actively secreted ECM proteins, including collagen VI within the interstitial matrix surrounding muscle fibers, and laminin gamma 1 in the basal lamina of newly formed fibers **(Fig. 4B)**. In the injected muscle, we specifically analyzed regions where human FAPs were present and compared them to adjacent regions lacking FAPs. Cross-sectional area (CSA) analysis revealed that muscle fibers in fibrotic FAPs-rich regions were significantly smaller than those in the rest of the TA **(Fig. 4C)**. Furthermore, quantification of endogenous Pax7+ cells showed a reduced number of muscle stem cells (MuSCs) in these fibrotic FAP-remodeled regions, suggesting that FAP-derived ECM may impair MuSCs quiescence and niche maintenance during regeneration **(Fig. 4D)**.

**Fig. 4.**
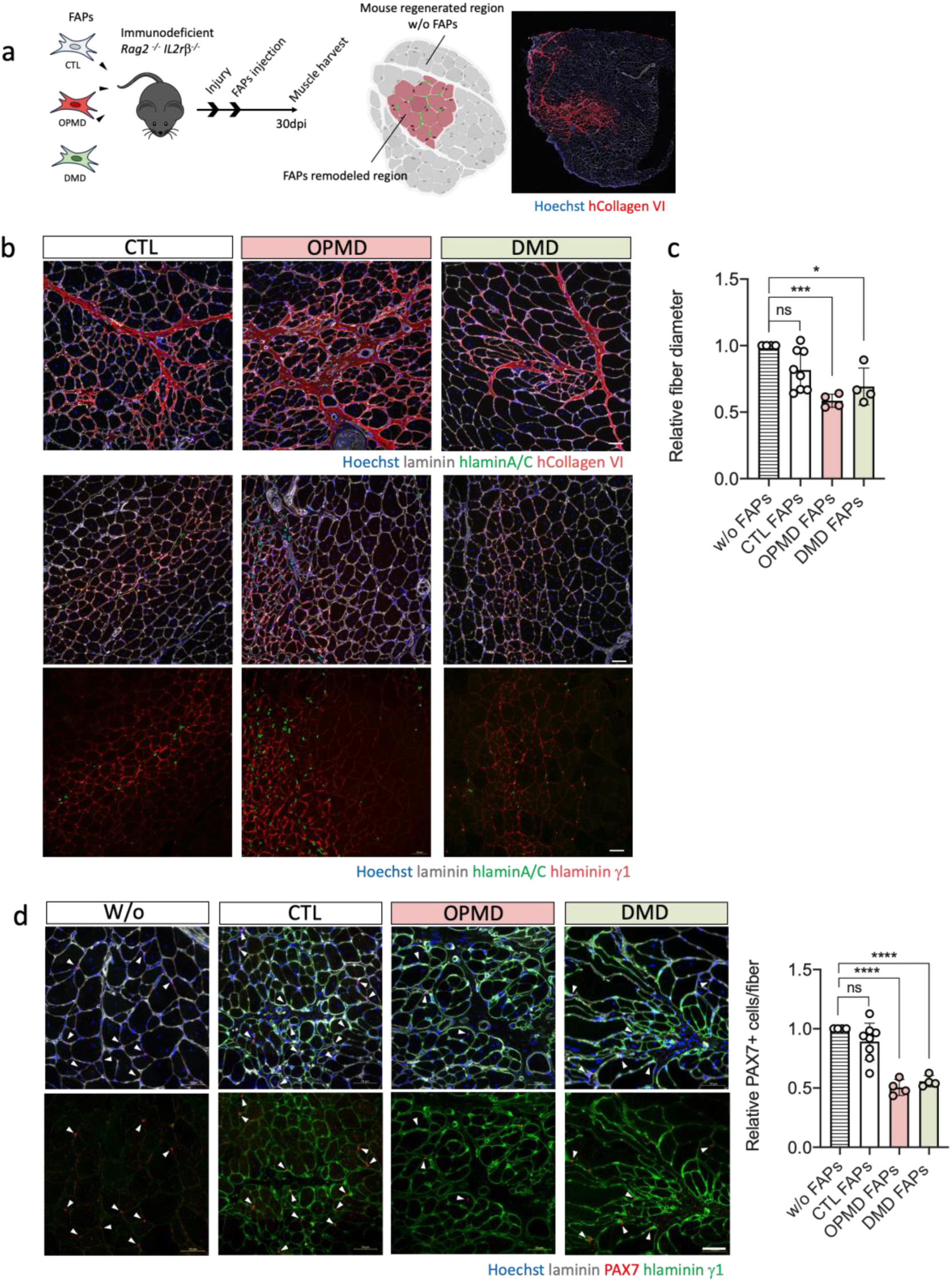
FAPs-driven ECM modifications impact muscle regeneration and stem cell niche maintenance. **(a**) Experimental set up of FAPs transplantation. FAPs from healthy CTL, OPMD or DMD were injected into the TA of immunodeficient mice after injury. TA were retrieved and analyzed 30 days post injections. For each analysis, the portion of the TA containing human cells (FAPs-remodeled regions) was compared to the regenerated mouse region without any human FAPs. **(b)** Immunostaining of laminin (white), Hoechst (blue) and human specific Collagen alpha-3(VI) chain (red) or human specific laminin gamma 1 (red). Scale bar: 50μm **(c)** Quantification of minimum ferret fiber diameter in the muscle region remodeled by FAPs normalized to the non-injected muscle area. **(d)** Immunostaining of laminin (white), Hoechst (blue), Pax7 (red) and human specific laminin gamma 1 (green). Arrows point to Pax7 positive cells. Scale bar: 50μm - Quantification of Pax7+ cells per fiber normalized to the non-injected muscle area. Data are presented as mean ±SD, with P-values obtained by ordinary one-way ANOVA test followed by Tukey’s multiple comparisons test (*P < 0.1 **P < 0.01, ***P < 0.001, ****P < 0.0001).

### Collagen VI plays a key role in myogenic fusion

Collagen VI was consistently upregulated in all three pathologies **(Fig. 3A)** and actively secreted by FAPs *in vitro* **(Fig. 3F**). Using human-specific primer sequences, we analysed the expression of several ECM candidates in FAPs injected into regenerating TA muscles of mice. Some mRNAs were significantly more highly expressed by disease-derived FAPs compared to controls. Among these, *Col6a1* and *Col6a3* were upregulated in FAPs from both pathologies in this regenerative context **(Fig. 5A)**. Moreover, public single-cell transcriptomic data show that *Col6a1, Col6a2* and *Col6a3* expressions are markedly higher in FAPs compared to other muscle-resident cell types **(Fig. 5B)**. To further investigate the impact of FAP-derived ECM on myogenic potential, we coated culture wells with increasing concentrations of collagen VI and assessed its effect on myogenic differentiation, as measured by fusion index *in vitro*. We found that higher concentrations of collagen VI significantly reduced the fusion ability of human muscle cells in a dose-dependent manner **(Fig. 5C).** To further directly assess the role of COLVI secreted by FAPs in the regulation of the fusion index, we silenced *COL6* expression specifically in FAPs isolated from OPMD patients using siRNA **(Fig. 5D-E)**. These siRNA-treated-FAPs were then co-cultured with myotubes for 3 days, and myogenic fusion was evaluated. The level of secreted COLVI expression, assessed by dot blot, was restored to the baseline level observed in wells without FAPs (**Fig**. **5E**). Notably, knockdown of *COL6* in FAPs (siCOL6) significantly improved the fusion index compared to co-cultures with FAPs transfected with a control siRNA (siCTL), indicating that FAP-derived COLVI contributes to the impaired myogenic fusion observed in disease conditions **(Fig. 5F)**.

**Fig. 5.**
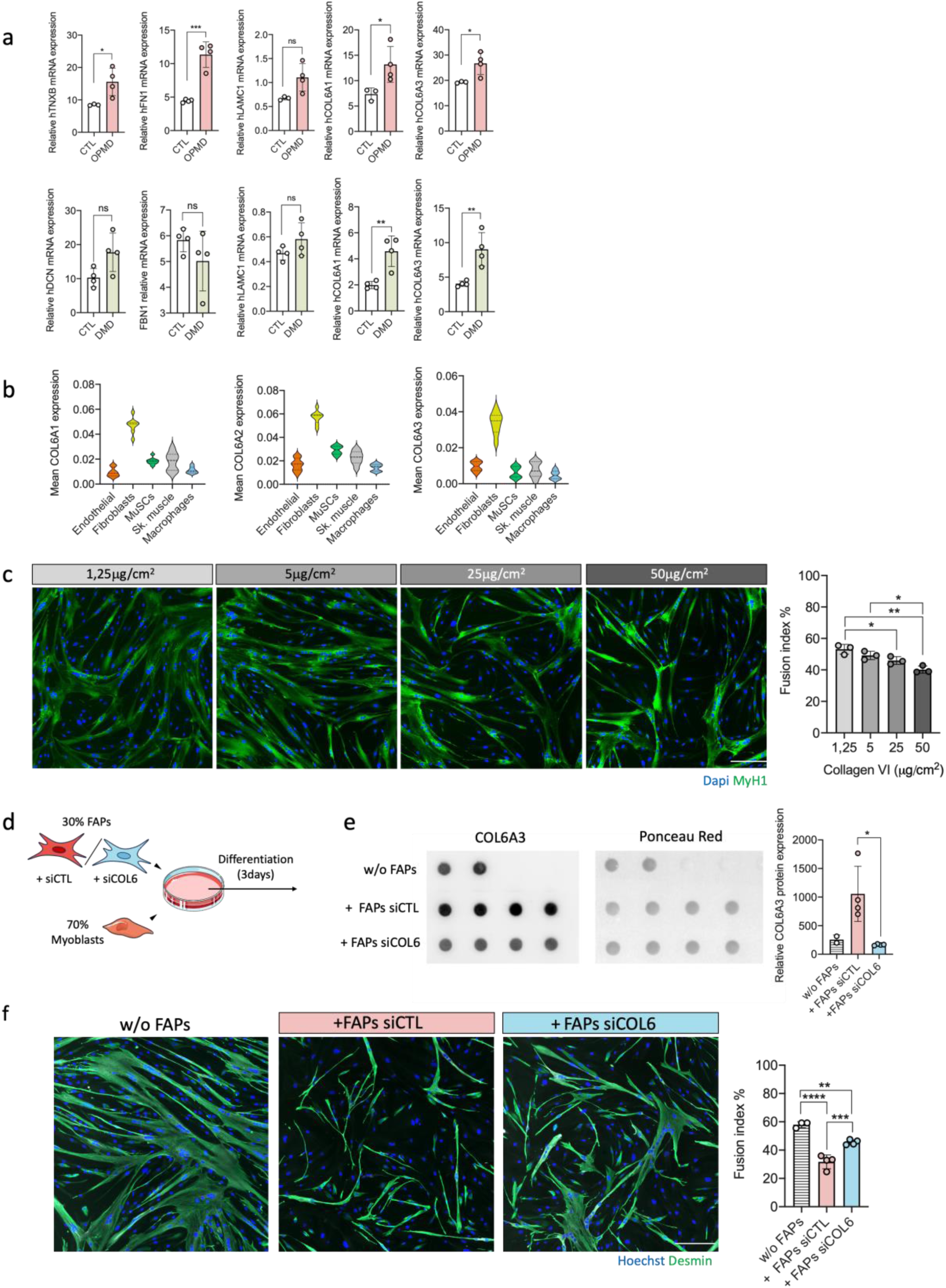
Collagen VI is involved in the impairment of myoblasts fusion *in vitro*. **(a)** RT-qPCR quantification of ECM genes normalized to the expression of human specific B2M **(b)** Expression of COL6A1, COL6A2 and COL6A3 in several resident cell types within skeletal muscle on sc-RNA-seq dataset extracted from De Micheli *et al* 2020 (GSE143704) **(c)** Immunostaining of myotubes (Myosin Heavy Chain, green; Hoechst, blue) after 5 days of differentiation on different concentration of collagen type VI coatings - Quantification of the fusion index. Scale bar: 200μm. **(d)** Schematic representation of co-culture experiment; Myoblasts were seeded with OPMD FAPs previously transfected for 24h with a siRNA against COL6A3 (siCOL6) or non-targeting siRNA (siCTL) and differentiated for 3 days **(e)** Quantification of Collagen VI alpha-3protein by dot blot on supernatants. Collagen signal was normalized on Ponceau red staining. **(f)** Immunostaining of myoblasts/FAPs co-cultures (Desmin, green; Hoechst, blue) after 3 days of differentiation - Quantification of the fusion index of myoblasts alone, or with FAPs transfected with COL6A3 siRNA or a non-targeting siRNA. Scale bar: 200μm

## Discussion

The ECM plays a central role in maintaining muscle architecture and regulating regeneration, yet its composition and remodeling dynamics in human myopathies remain poorly defined. While fibrosis, defined as an excessive deposition of ECM, is a common feature across many muscular dystrophies, most studies have focused on inflammatory or myogenic aspects, leaving the matrisome comparatively understudied. How ECM remodeling differs—or converges—across distinct fibrotic muscle diseases has not been systematically addressed. Here, we provide a comparative analysis of ECM composition in DMD, OPMD, and IBM. By integrating proteomic profiling with functional assays, we identify both shared and disease-specific ECM signatures and demonstrate that FAP-mediated ECM remodeling contributes to impaired muscle regeneration through disruption of the stem cell niche, allowing fibrotic tissue to develop within the defective regenerating muscle.

A specific characteristic of the ECM is its capacity to undergo dynamic remodeling, balancing synthesis and degradation in a tightly regulated manner ^16^. In all three muscle diseases – DMD, OPMD and IBM – we observed a common shift toward impaired ECM turnover, primarily driven by increased expression of protease inhibitors, as illustrated by the examples described below. Protease inhibitors such as A2M and SERPINA1 have been highlighted in DMD ^17^ and our data also reveal elevated levels of other serine protease inhibitors like SERPINC1 in both DMD and OPMD. This pattern of broad protease inhibition suggests a common mechanism limiting ECM degradation in these muscle diseases. Additionally, the plasminogen (PLG) system, a key player in ECM remodeling through fibrin and fibronectin degradation and activation of protease ^18^, was elevated in OPMD and IBM, and detected in DMD compared to controls. Together, these changes likely contribute to pathological ECM accumulation and fibrosis, reflecting a shared disruption of ECM homeostasis across all three diseases.

This impaired ECM turnover also intersects with inflammatory processes, which are a hallmark of DMD, IBM, and OPMD. In these diseases, the accumulation of ECM-derived DAMPs (damage-associated molecular patterns) and inflammatory mediators contributes to chronic inflammation and fibrosis. Notably, increased annexins (ANXA1, ANXA2) point to active membrane repair mechanisms and a potential role in sustaining fibrotic signaling. Moreover, the presence of pro-inflammatory molecules such as fibrinogen—already elevated in the ECM of DMD and OPMD—can activate macrophages and promote TGFβ signaling, further exacerbating tissue fibrosis ^19^.

Both OPMD and IBM are late-onset disorders characterized by mitochondrial dysfunction ^4,20^ and protein inclusions. Before filtering our proteomic data to focus on the matrisome, we observed upregulation of several mitochondrial enzymes (ACO2, CS, UQCRC1) and molecular chaperones (e.g. HSPA9, HSPA1A) indicative of mitochondrial stress and activation of protein quality control mechanisms. These findings are consistent with the cellular stress responses typically observed in these conditions. Although OPMD and IBM differ in their genetic and pathological origins— PABPN1 mutations in OPMD versus the inflammatory and degenerative processes in IBM—they share some overlapping clinical features. Notably, both disorders commonly present with dysphagia, reflecting their preferential involvement of the cricopharyngeal muscle. Following matrisome-specific filtering, our analysis revealed that OPMD and IBM display distinct ECM remodeling signatures, including differences in collagen composition and the expression of ECM-regulatory proteins such as fibrillin-1 (FBN1). These alterations are particularly relevant given the role of the ECM as a reservoir for bioactive molecules, including TGFβ. Differential levels of FBN1 across diseases suggest disease-specific regulation of TGFβ activation: reduced FBN1 in DMD may increase TGFβ bioavailability, while elevated FBN1 in IBM could sequester or modulate its activation through ECM structural remodeling ^21^. These findings reinforce the idea that altered ECM composition can directly influence profibrotic signaling and contribute to disease progression.

DMD’s pathogenesis results from dystrophin deficiency leading to sarcolemma destabilization, fiber damage and chronic inflammation ^22^. Our global proteomic data showed increased expression of cytoskeletal proteins (such as vimentin and various myosin isoforms) and glycolytic enzymes (GPI, LDHA, PKM), reflecting cytoskeletal reorganization and metabolic adaptation to ongoing muscle regeneration. Following matrisome-specific filtering, we observed that the fibrotic response in DMD involves extensive ECM remodeling. This includes upregulation of protease inhibitors (A2M, SERPINA1), increased collagen deposition, and elevated levels of pro-inflammatory mediators like fibrinogen ^17,23^, which contribute to the amplification of fibrotic signaling and tissue scarring. In contrast, IBM displayed a distinct ECM remodeling profile, particularly marked by the accumulation of small leucine-rich proteoglycans (SLRPs) such as decorin and lumican. These proteins are key regulators of collagen fibril stability and play an important role in modulating muscle repair dynamics.

ECM structure and stiffness are also shaped by its proteoglycan content. Several small leucine-rich proteoglycans (SLRPs)—including ASPN, BGN, DCN, LUM, OGN, and PRELP—bind collagen fibrils to protect them from collagenase cleavage ^24,25^. Notably, DMD samples showed reduced levels of ASPN, LUM, OGN, and PRELP, with DCN increased and BGN uniquely enriched in DMD samples. As SLRPs compete for collagen-binding sites (e.g., ASPN and DCN; ^26^), these imbalances could destabilize collagen architecture and alter matrix stiffness. ECM stiffness directly influences muscle cell fate: Increased stiffness has been shown to modulate FAP differentiation ^27^ and impair satellite cell activation ^28,29^. Thus, shared ECM remodeling defects in DMD, OPMD, and IBM may converge on a common mechanism: reduced matrix turnover coupled with altered stiffness, reinforcing fibrosis, impairing stem cell activation and disrupting regenerative signaling.

Beyond its impact on fibrosis, ECM composition and stiffness also directly affect muscle stem cell (MuSC) behavior, particularly within the specialized microenvironment of the basal lamina, which is rich in collagens and laminins ^30^. This niche governs MuSC quiescence, activation, and self-renewal, making it essential for muscle homeostasis and regeneration ^31^. Specific ECM components have been shown to exert opposing effects on MuSC fate: some, such as laminin 211, promote quiescence ^8^, while others, like laminins containing the α5 chain, support proliferation ^32^. Our findings demonstrate that disease-derived FAPs significantly remodel the ECM during regeneration, with consequences for MuSC maintenance. When human FAPs from fibrotic muscle biopsies were transplanted into regenerating mouse muscle, they integrated into the interstitial space and actively secreted ECM proteins, including collagen VI and laminin subunits. Importantly, regions enriched in fibrotic FAPs displayed smaller muscle fibers and a reduced number of Pax7+ MuSCs, suggesting that excessive or specific ECM deposition impairs MuSC quiescence, niche integrity and on-going regeneration. In the proteomic data from all three diseases, we observed elevated levels of basal lamina components such as LAMA2 (merosin), LAMB2, LAMC1 and COL4A1. These alterations may disrupt MuSC positioning and niche integrity, contributing to impaired MuSC behavior and defective regeneration observed in muscle biopsies from OPMD ^33^, DMD ^34^ and IBM patients ^35^.

Among the ECM components identified in our study, collagen VI stands out as a key player. It was consistently increased across all disease conditions and secreted by FAPs both *in vitro* and *in vivo*. Functionally, increased collagen VI impairs myogenic differentiation, as shown by reduced fusion indices in myoblast cultures exposed to high COLVI levels. Previous studies carried out in mice highlight the finely regulated role of collagen VI in modulating the behavior of satellite cells. A complete deficiency of collagen VI impairs MuSC self-renewal following injury ^36^ emphasizing its importance in maintaining a functional niche and promoting proper muscle regeneration. Indeed, supplementation with collagen VI—via mesenchymal stem cells expressing collagen VI—enhances muscle regeneration and maturation in Col6ko mouse models ^37^. Elevated levels of collagen VI, by inhibiting the canonical Wnt pathway (which acts as a soluble ligand), delay early myogenic differentiation ^38^. Our experiments using cultured human myoblasts appear to reflect a similar pattern. Recently Mohassel et al ^39^ showed that during murine muscle regeneration collagen VI-deficient matrix is not able to regulate the dynamic TGF-β bioavailability, impairing this regulatory mechanism.

Taken together, our data reveal that FAP-mediated ECM remodeling—particularly the aberrant deposition of collagens and laminins—profoundly alters the MuSC niche, impairing stem cell positioning, quiescence, and regenerative potential. These findings reposition the ECM not merely as a passive structural scaffold but as an active, disease-modified microenvironmental cue with instructive roles in stem cell fate decisions. By delineating both shared and disease-specific ECM alterations, our study provides a conceptual framework for understanding how niche dysfunction contributes to regeneration failure in fibrotic myopathies. We uncovered Collagen 6 as a conserved pro-fibrotic signal disrupting muscle regeneration across distinct human myopathies. Our findings further underscore the complexity of ECM remodeling in fibrotic muscle diseases: while mechanisms such as increased collagen deposition and protease inhibition are commonly observed, distinct disease-specific ECM signatures reveal divergent pathogenic processes. Importantly, these signatures may offer novel targets for therapeutic strategies aimed at restoring MuSC function and muscle homeostasis. Future studies should now focus on mechanistically linking specific ECM alterations to MuSC dysfunction and evaluating targeted interventions in relevant disease models.

## Materials and Methods

### Muscle biopsy samples

Human muscle samples from healthy controls (CTL), oculopharyngeal muscular dystrophy (OPMD), Duchenne muscular dystrophy (DMD) and inclusion body myositis (IBM) patients (see Table 5) were obtained anonymously via the Myobank-AFM, affiliated with EurobioBank, in accordance with European recommendations and French legislation (authorization AC-2019-3502).

**Table 5.** Human biopsies used in the study.

| Status | Muscle | Number |
| --- | --- | --- |
| CTL | Cricopharyngeal | 4 |
|  | Dorsal | 1 |
|  | Deltoid | 1 |
|  | Paraspinal | 5 |
|  | Quadriceps | 6 |
|  | Tensor fascia lata | 1 |
|  | Sternocleidomastoid | 9 |
| OPMD | Cricopharyngeal | 13 |
| IBM | Cricopharyngeal | 11 |
| DMD | Paraspinal | 8 |
|  | Quadriceps | 1 |
|  | Spinal | 1 |

### Muscle histology

Fresh human muscle biopsy samples (OPMD, IBM, DMD and CTL) were mounted on tragacanth gum (6% in water; Sigma-Aldrich) placed on a cork support and snap frozen in liquid nitrogen-cooled isopentane. Transverse serial frozen cryosections (5μm thick) obtained with a cryostat (Leica CM1850) were processed for staining. Sirius red (SR) coloration was obtained by first fixing the sections with 4% paraformaldehyde (PFA) followed by an incubation in a 0,3% solution of Sirius red at room temperature (RT) for 1h. Sections were imaged by light microscopy (Leica DMR microscope equipped with a Nikon DS-Ri1 camera). The percentage of fibrosis within each biopsy was assessed on SR stained cryosections from 3 to 5 fields per sample, using ImageJ software, and expressed as a percentage of the total area analyzed. Cryosections were used for subsequent proteomic and immunofluorescence analyses.

### Proteomic sample preparation

The preparation of tissue specimens and subsequent analysis of extracted proteins by bottom-up proteomics was performed as previously described ^40^. Frozen cryosections (for a total amount of 250 μm) from human muscle biopsy samples (OPMD, IBM, DMD and CTL) were homogenized using lysis buffer containing 4% (w/v) sodium dodecyl sulfate, 0.1 M dithiothreitol and 100 mM Tris-Cl, pH 7.6 and a protease inhibitor cocktail (Thermo Scientific #A32953). Homogenization was carried out using a handheld homogenizer (Kimble Chase, Rockwood, TN, USA), followed by a brief water bath sonication. Samples were then heated for 3 minutes (min) at 95 °C and centrifuged at 16,000*g* for 5 min. The protein-containing supernatant was extracted and used for subsequent proteomic studies. The Pierce 660 nm Protein Assay system was used to determine protein concentration. Extracted proteins were further processed for mass spectrometric analysis. Samples were mixed with 8 M urea, 0.1 M Tris pH 8.9 in Vivacon 500 spin filter units and centrifuged at 14,000*g* for 15 min. Samples were processed with Filter-aided sample preparation (FASP) method ^41^, buffer switching and protein trypsination prior to mass spectrometric peptide analysis.

### Mass spectrometry and proteomic data analysis

Protein samples were analysed using a Q-Exactive mass spectrometer (Thermo Scientific) coupled to a Dionex RSLCnano (Thermo Scientific). Peptides were separated using a 2–40% gradient of acetonitrile (A: 0.1% FA, B: 80% acetonitrile, 0.1% FA) over 65 min at a flow rate of 250 nl/min. The Q Exactive was operated in the data dependent mode, collecting a full MS scan from 300–1650 m/z at 70 K resolution and an AGC target of 1e6. The 10 most abundant ions per scan were selected for MS/MS at 17.5 K resolution and AGC target of 1e5 and intensity threshold of 1 K. Maximum fill times were 10 ms and 100 ms for MS and MS/MS scans respectively with a dynamic exclusion of 60 s. Samples were analysed using 25 NCE (normalized collisional energy) with 20% stepped energy ^42^. For quantitative analysis, samples were evaluated using MaxQuant (v1.5.2.8) and the Andromeda search engine to identify the detected features against the UniProtKB/SwissProt database (Homo sapiens). The following search parameters were used: (i) first search peptide tolerance of 20 ppm, (ii) main search peptide tolerance of 4.5 ppm, (iii) cysteine carbamidomethylation set as a fixed modification, (iv) methionine oxidation set as a variable modification, (v) a maximum of two missed cleavage sites and (vi) a minimum peptide length of seven amino acids. The false discovery rate (FDR) was set to 1% for both peptides and proteins using a target-decoy approach. Perseus v.1.5.6.0 was used for data analysis, processing, and visualization ^43^.

### Proteomic Data Analysis

Cricopharyngeal (CP) muscles from OPMD (n=3) and IBM patients, (n=4) were compared to CP muscle from age-matched healthy subjects (n=4). DMD paraspinal (PS) muscles (n=4), were compared to PS from age-matched healthy subjects (see table 6). Enriched terms for biological processes from the list of differentially expressed proteins were obtained using g: Profiler ^44^. Differentially expressed genes were analysed with the Retrieve/ID mapping tool from the UniProt database to obtain the corresponding reviewed protein sequences. These sequences were then used to obtain information about the presence of signal peptides, transmembrane domains, and unconventional protein secretion using OutCyte v1.022 prediction tools. Lists of proteins have been filtered using the list from the Matrisome project (http://matrisomeproject.mit.edu) ^14^ to annotate them into matrisome protein groups.

**Table 6.**
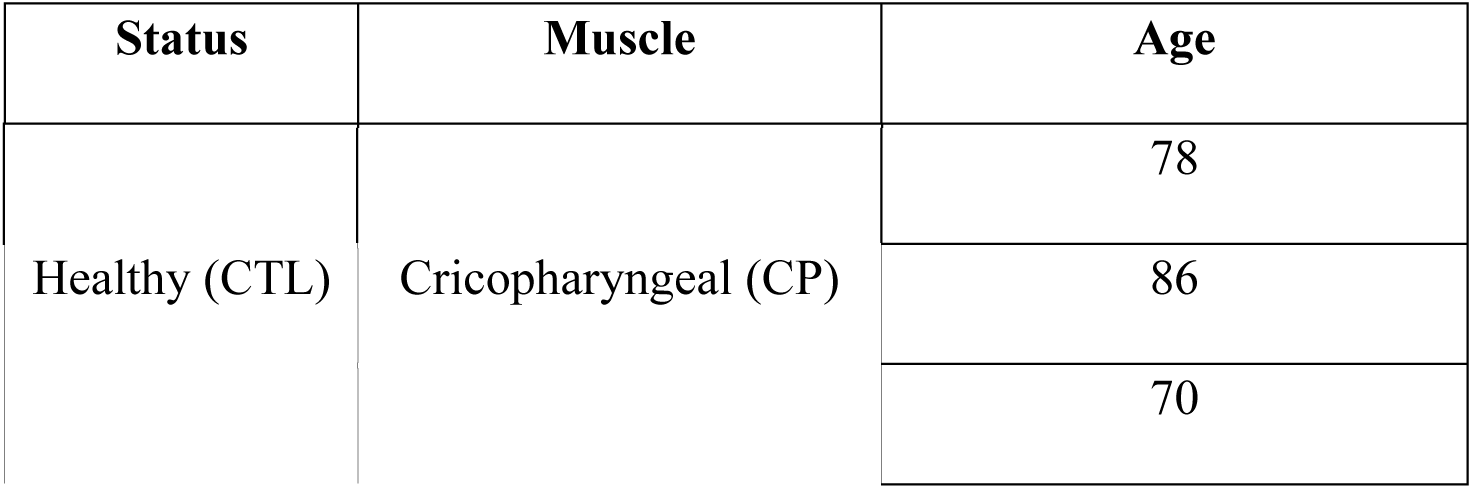
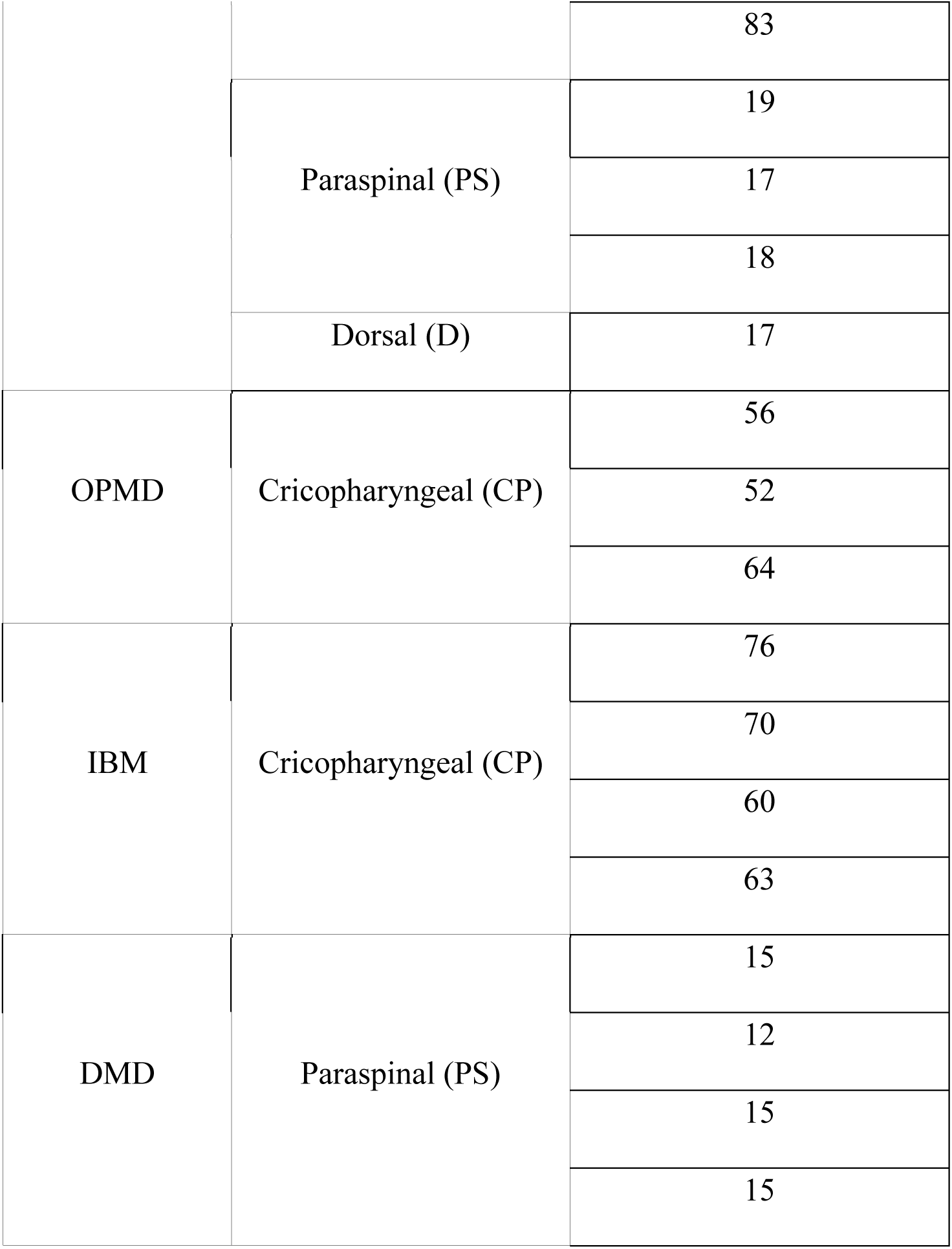
Human biopsies used for the LC-MS/MS analysis.

### Cell isolation, culture and ECM production

Human cells from muscle samples were isolated as described previously ^45^: briefly, muscle biopsy samples were minced, and explants plated onto dishes coated with fetal bovine serum (FBS) (Invitrogen). Isolated cells were cultured at 37°C in growth medium (GM) consisting of 199 medium (Life Technologies) and Dulbecco’s modified Eagle’s medium (DMEM, Life Technologies) at a 1:4 volume ratio supplemented with 20% FBS, 25 μg/mL fetuin (Life Technologies), 0.5 ng/ mL bFGF (Life Technologies), 5 ng/mL EGF (Life Technologies), 5 μg/mL insulin (Sigma-Aldrich), and 50 μg/mL gentamycin (Life Technologies) in a humid atmosphere containing 5% CO_2_. Myoblasts and FAPs were isolated by using magnetic activated cell sorting (MACS) with CD56 antibody-coupled microbeads (130-050-401, Miltenyi Biotec) according to the manufacturer’s instructions. After separation of the two populations; CD56+ myoblasts and CD56-FAPs; the purity of each population was monitored by immunocytochemistry after fixation for 10 min with pure ethanol using an antibody against desmin (clone D33, Dako), which is exclusively expressed in myoblasts. At least 500 cells per condition were counted. FAPs used in subsequent analysis were always <1.5% desmin-positive and myoblasts >90% desmin-positive. Lifespan and proliferative status of cells used in this study have been carefully monitored to make sure cells do not reach pre-senescence for any of the conditions. For production of ECM, FAPs were plated at 15.10^e^3/cm^2^ on non-coated glass coverslips in 4-well plates and the GM was changed every 3 days for 14 days.

### In vivo FAPs transplantation

This study was carried out in strict accordance with the legal regulations in France and according to the ethical guidelines for animal research of the European Union. The protocol was approved and delivered by the French Ministry of Higher Education and Scientific Research (number: 2021072217421004_v4).

Rag2 -/- Il2rb -/- immunodeficient mice were used as recipients of transplanted human FAPs. Mice were anaesthetized by an intraperitoneal injection of 80 mg/kg ketamine hydrochloride and 10 mg/kg xylazine. The *tibialis anterior* (TA) of the mice were subjected to damage in order to trigger regeneration; for OPMD FAPs injections, TAs were subjected to three freeze lesion cycles as previously described ^46^, then 15 μL of 1.4 × 10^5^ FAPs in PBS were injected immediately after cryodamage and 4 and 8 days later. For DMD FAPs injections, TAs were injected with 20μl of notexin 20μg/mL and 1 × 10^5^ cells in 15ul of PBS were injected the next day. Mice were euthanatized by cervical dislocation 4 weeks after the first injection, and the TAs were collected, snap frozen in isopentane, and stored at −80°C for further analyses. Human specific antibodies were used to distinguish human cells within mice TAs. For fiber diameter and Pax7 positive cells number analysis, areas containing human cells were compared to areas without human cells in the same mouse TA.

### Co-culture and collagen coating experiments

For co-culture experiments, myoblasts and FAPs were seeded together at a 70%/30% ratio and a final density of 21 000 cells/cm^2^ in μ-Slide 8 Well (80826, Ibidi) plates. Cells were differentiated by shifting growth medium to a serum-free medium consisting of DMEM and 50 μg/mL gentamycin. After 3 or 5 days of differentiation, cells were fixed with PFA 4% and used for subsequent immunostaining experiments.

For collagen treatment, collagen 6 (Rockland, 009-001-108) was diluted in 0,1% acetic acid and coated on μ-Slide 8 Well (80826, Ibidi) plates from 1,25 to 50 μg/cm^2^. Coating was left at 37°C during 1h before washing with PBS and seeding human myoblasts from healthy individuals. Cells were differentiated the next day and fixed with PFA 4% after 5 days of differentiation.

### siRNA transfection

Primary FAPs from OPMD patients were seeded at a confluence of 80% in 48-well plate. The next day, cells were transfected with COL6A3 siRNAs (ON-TARGETplus siRNA human COL6A3 SMARTPool, L-003646-00, Dharmacon) at a final concentration of 30nM, using Lipofectamine RNAiMAX reagent (10601435, Invitrogen) according to the manufacturer’s instructions, in growth medium. ON-TARGETplus Non targeting pool (D-001810-10) was used as a negative control. The next day, FAPs were detached, counted and seeded in coculture with myoblasts as described above. 3 days after differentiation, supernatant was collected and cells were fixed with PFA 4% for subsequent analysis.

### Dot blot analysis

Supernatant from co-culture experiments were collected, and centrifuged at 300G for 10 min to remove dead cells. Dot blot apparatus (Bio-Dot 1706545, Biorad) was used following manufacturer instructions: 40μl of each supernatant was transferred onto a nitrocellulose membrane by micro-filtration. Membrane was stained with Ponceau red for total protein quantification. After rinsing, membrane was blocked in Tris-buffered saline with Tween 20 (TBST) containing 5% milk for 30 min before incubation with COL6A3 antibody (MAB 1944) overnight. Membrane was incubated with appropriate secondary antibody in TBST for 45 minutes before revelation using the Clarity Max ECL substrate (1705062, Biorad) on a ChemiDoc MP imaging system (Biorad). Protein quantification was performed by normalizing Collagen VI signal to total protein using an imageJ macro (dx.doi.org/10.17504/protocols.io.7vghn3w).

### Quantitative PCR

RNA from frozen mouse muscle sections was extracted using Trizol reagent (Invitrogen) according to the manufacturer’s instructions. RNA was reverse transcribed using M-MLV (Invitrogen) according to the manufacturer’s instructions. Quantitative polymerase chain reaction (qPCR) was carried out using SYBR Green Mastermix (Roche Applied Science) in a LightCycler 480 Real-Time PCR System (Roche, Applied Science) or a Applied Biosystems QuantStudio 6 Pro (Thermofisher scientific) with the following cycling protocol: 8 min at 95°C; followed by 50 cycles at 95°C for 15 s (s), 60°C for 15 s and 72°C for 15 s, and a final step consisting of 5 s at 95°C and 1 min at 65°C. The specificity of the PCR product was evaluated by melting curve analysis using the following program: 65°C to 97°C with a 0.11°C/s increase. To analyze the gene expression of injected human cells, only human specific primers (see table 7) were used and gene expression levels were normalized to human RPLP0 or human B2M expression and quantified with the 2–ΔΔCt method. The human specificity of the primers was attested using a non-injected TA muscle as a negative control for each qPCR. The primer sequences used in this study are listed in the table below.

**Table 7.**
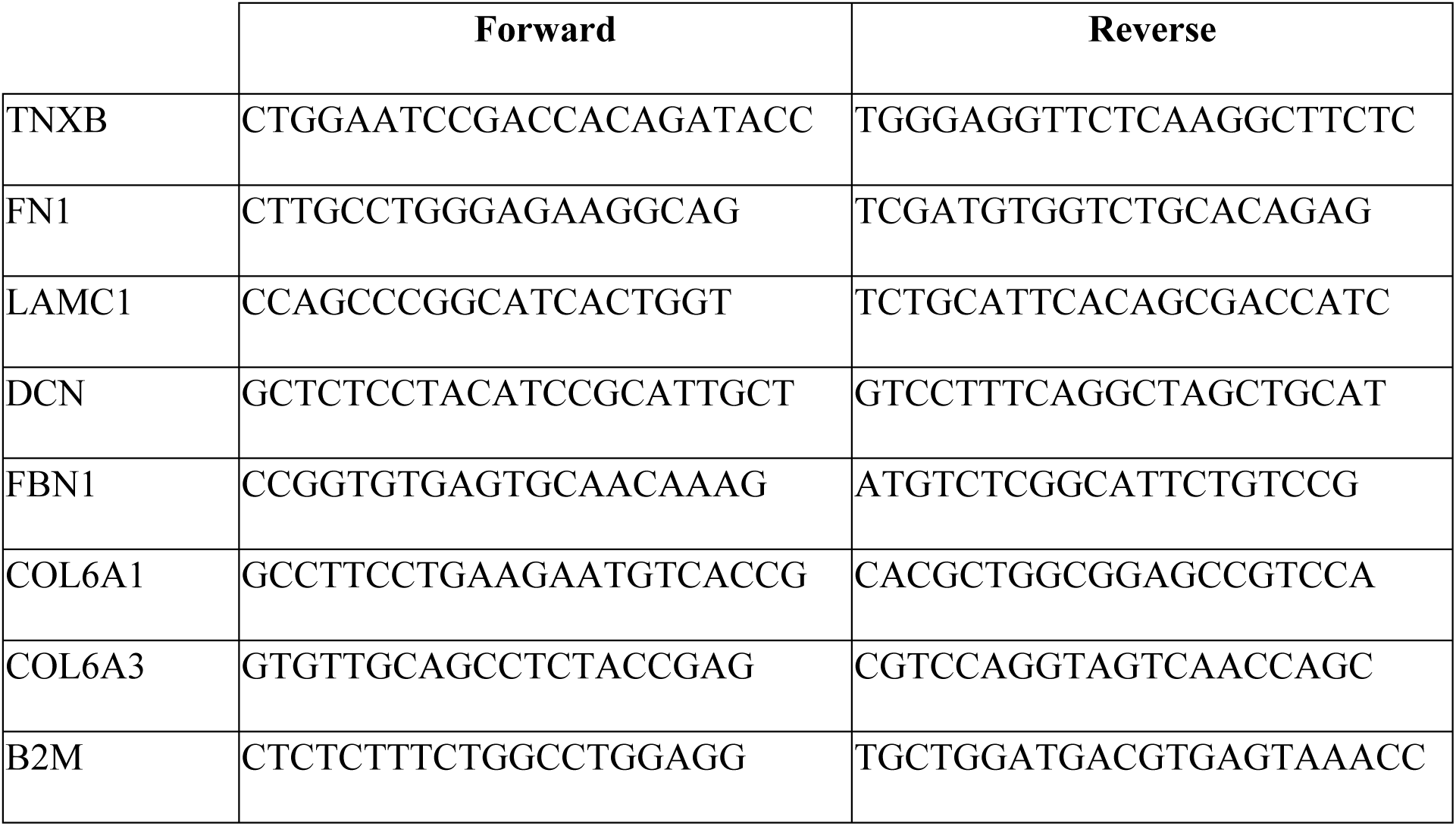
Primers used for quantitative PCR.

### Immunofluorescence imaging

Immunostaining was performed either on 5μm thick muscle biopsy sections or on FAPs *in vitro*, fixed in 4% paraformaldehyde for 10 min at room temperature (RT) and incubated with PBS containing 2% FBS and 0,2% triton for 30 min at RT. Incubation with primary antibodies (see table 8) was performed at RT for 1 hour (h) then cryosections or fixed cells were incubated with appropriate fluorescent secondary antibodies (Life technologies) at 1/400 at RT for 45 min. Nuclei were stained with Hoechst. Widefield fluorescence images were taken with an Axio Observer 7 microscope (Zeiss) equipped with a motorized stage coupled to an Orca Flash 4 Camera (Hamamatsu) and driven by the Zen software (Zeiss).

**Table 8.**
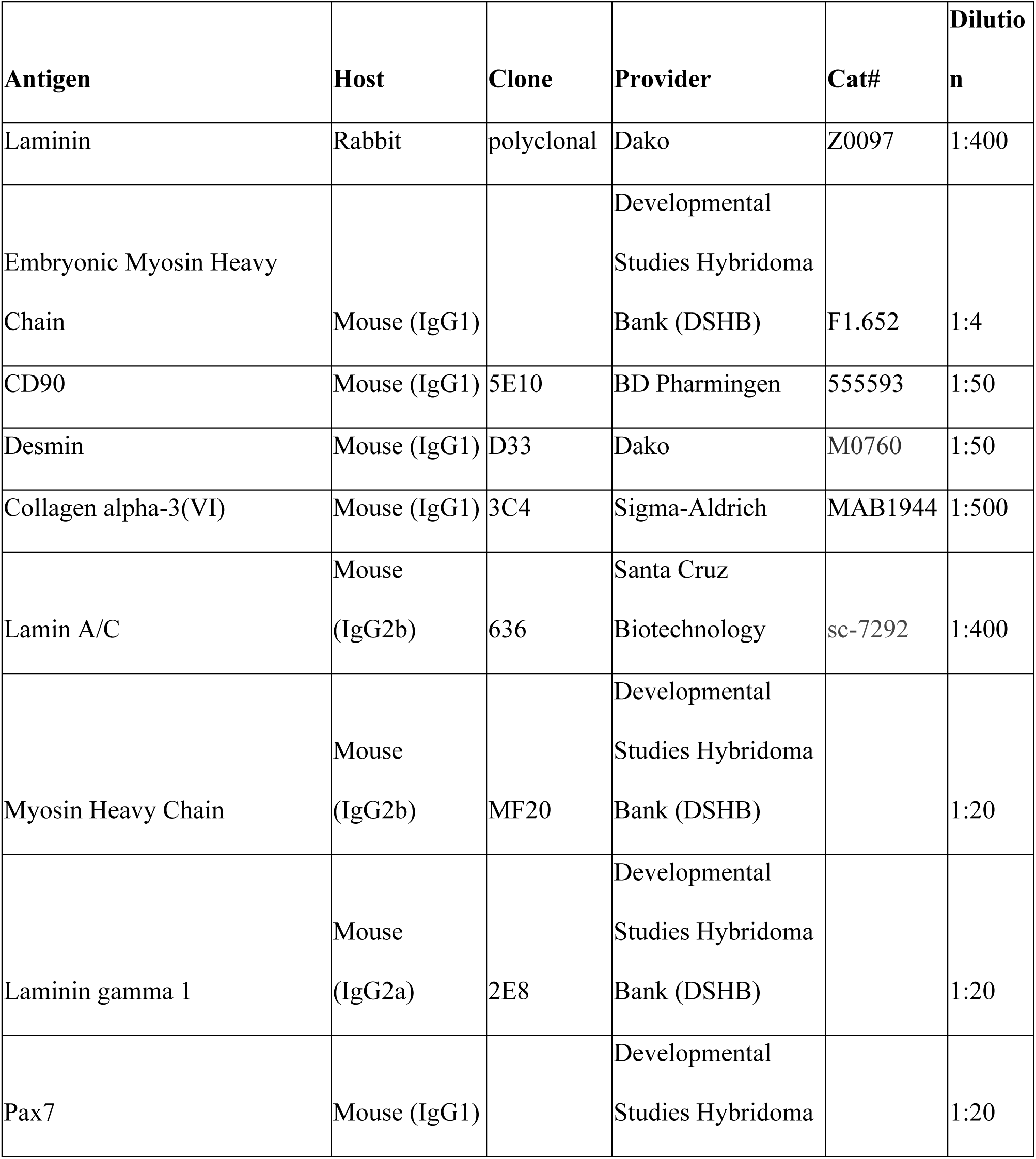
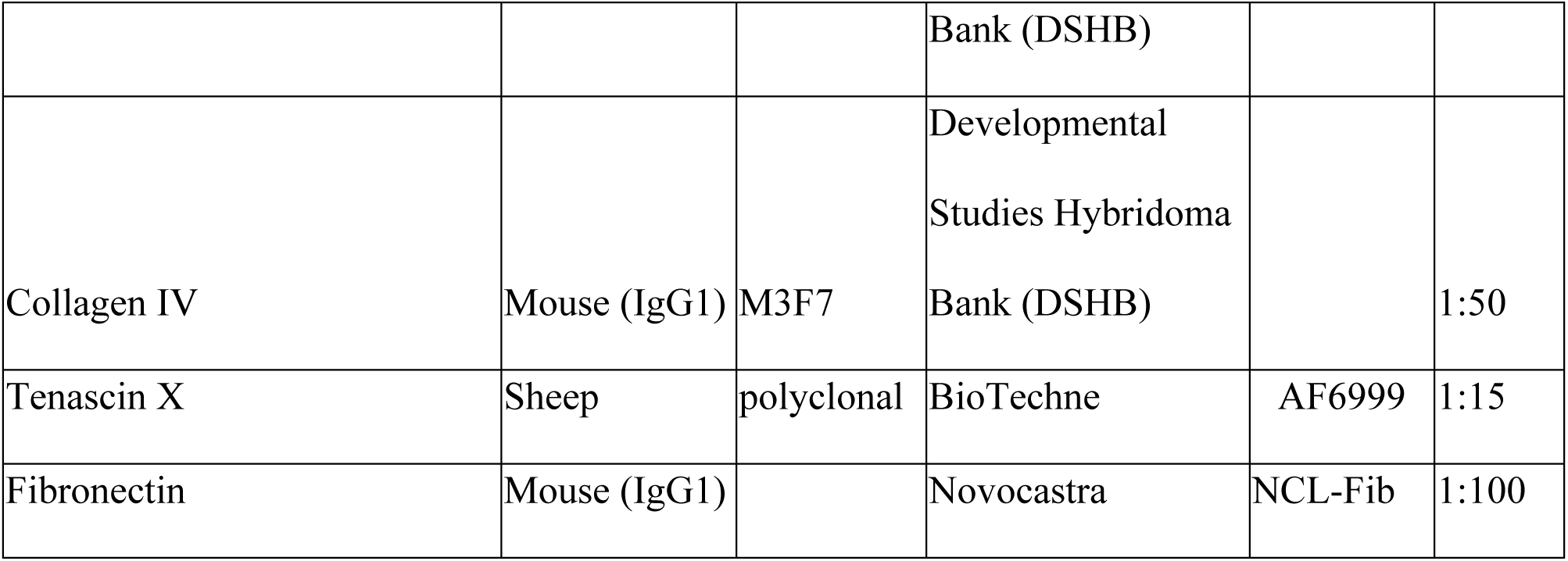
Antibodies used for immunofluorescence microscopy

### Statistics

Data are expressed as the mean ± SD. Statistical significance was assessed by ordinary or RM one-way ANOVA followed by Tukey’s multiple comparisons test or student T tests. All statistical analyses were performed using GraphPad Prism (version 8.0, GraphPad Software Inc.). Differences were considered to be significant at *P < 0.05, **P < 0.01, ***P < 0.001, and ****P < 0.0001.

### Data and materials availability

All data needed to evaluate the conclusions in the paper are present in the paper and/or the supplementary Materials. The mass spectrometry proteomics data have been deposited to the OSF website (https://osf.io/uejpv/?view_only=661b12b732354b35a3a7f9e50afcfd56).

## Acknowledgments

The authors would like to dedicate this work to the memory of their colleague and friend Didier Montarras, who spent his career studying skeletal muscle biology and whose passion and discoveries deeply shaped the field. We thank the clinicians and Stéphane Vasseur and Maud Chapart from the MyoBank for providing muscle surgical wastes from patients with informed consent. We thank the MyoIMAGE facility for imaging support. We thank Dimitris Kourtzas for his help on the dot blot analysis.

## Author contributions

L.M., C.T., and E.N. contributed to the conceptualization.

L.M., M.B., S.G., P.D., M.K., and V.A. contributed to the methodology.

L.M., M.B., J.D., S.G., P.D., E.N., and A.B. carried out the investigation.

C.T., E.N., and K.O. provided supervision.

L.M., E.N., and C.T. wrote the original draft.

L.M., M.B., J.D., M.K., S.G., P.D., A.B., V.A., S.P., J.L.G., G.B.B., V.M., K.O., C.T., and E.N. reviewed and edited the manuscript.

## Funding

This work was supported by funding from the Institut National de la Santé Et de la Recherche Médicale (INSERM), Sorbonne Université, Association Institut de Myologie, Fondation pour la Recherche Médicale FRM (EQUIPE FRM EQU201903007784), Agence Nationale pour la Recherche (ANR, «EpiMUSE» grant # ANR-22-CE14-0068; “FibroDys” grant #ANR-24-CE14-5209; “CPJ” grant #ANR-24-CPJ1-0122-01), Kathleen Lonsdale Institute for Human Health Research at Maynooth University, Science Foundation Ireland Infrastructure Award SFI-12/RI/2346/3, Research Ireland and Campus France (Ulysses/2024/5)

## Competing interests

Authors declare that they have no competing interests

---

**Table 1A.**
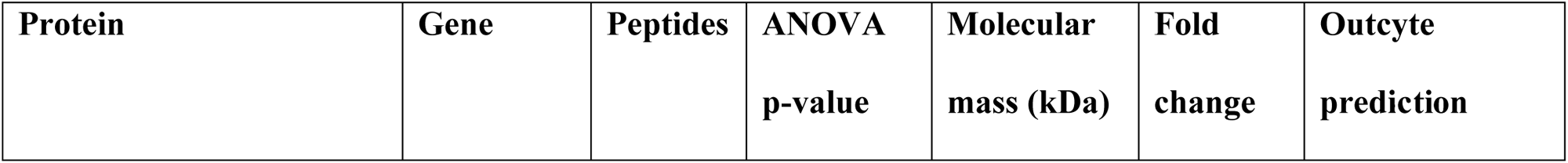
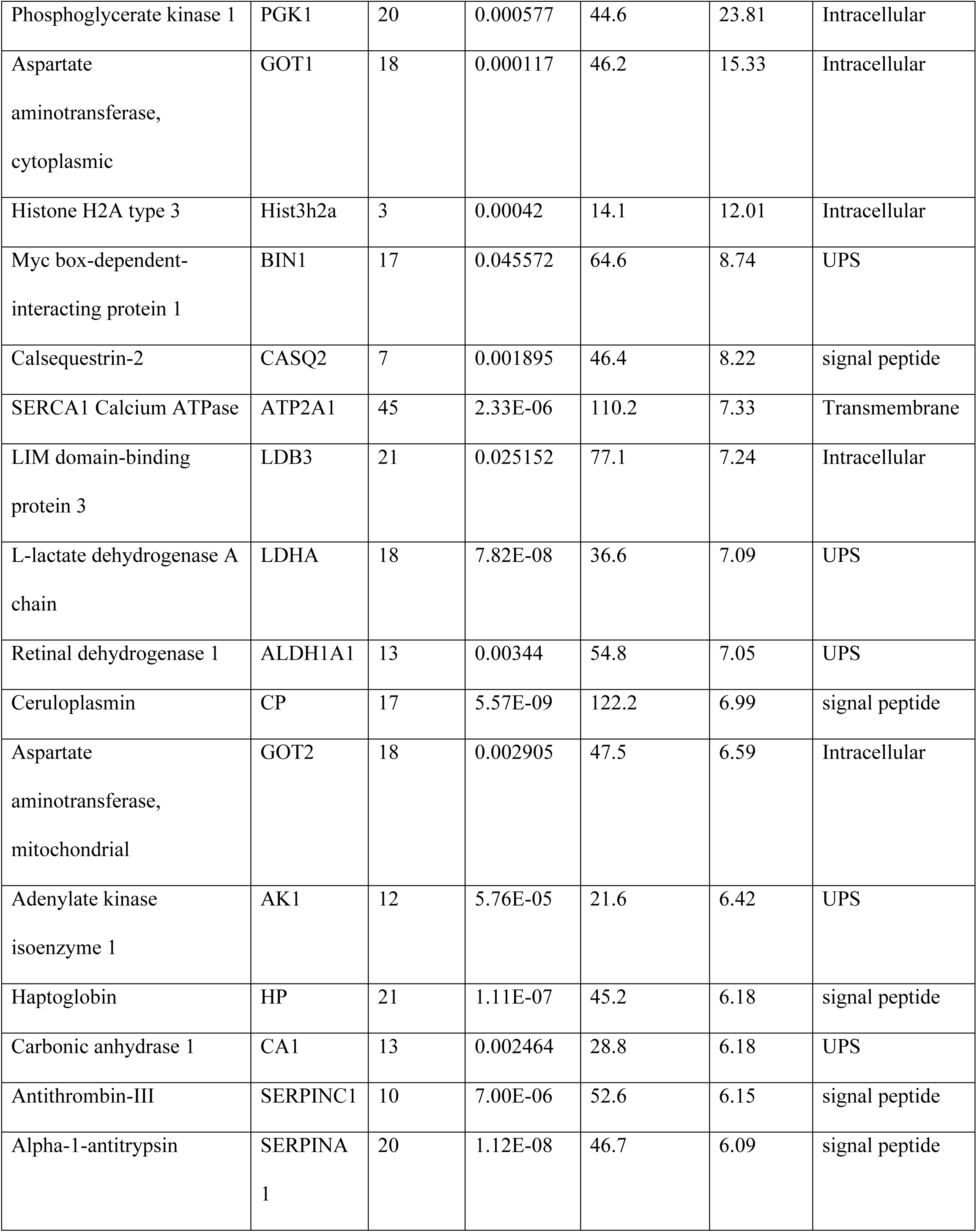
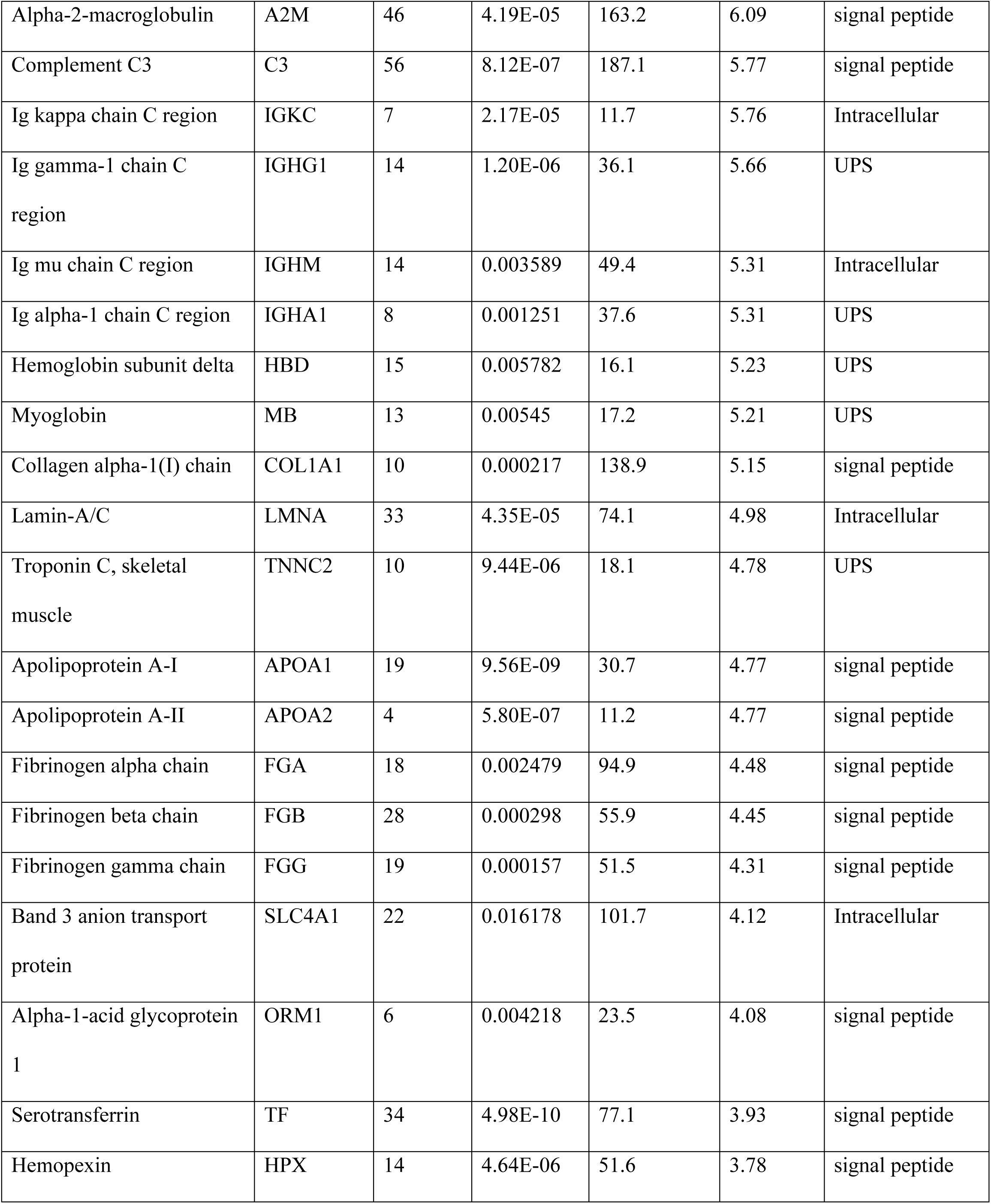
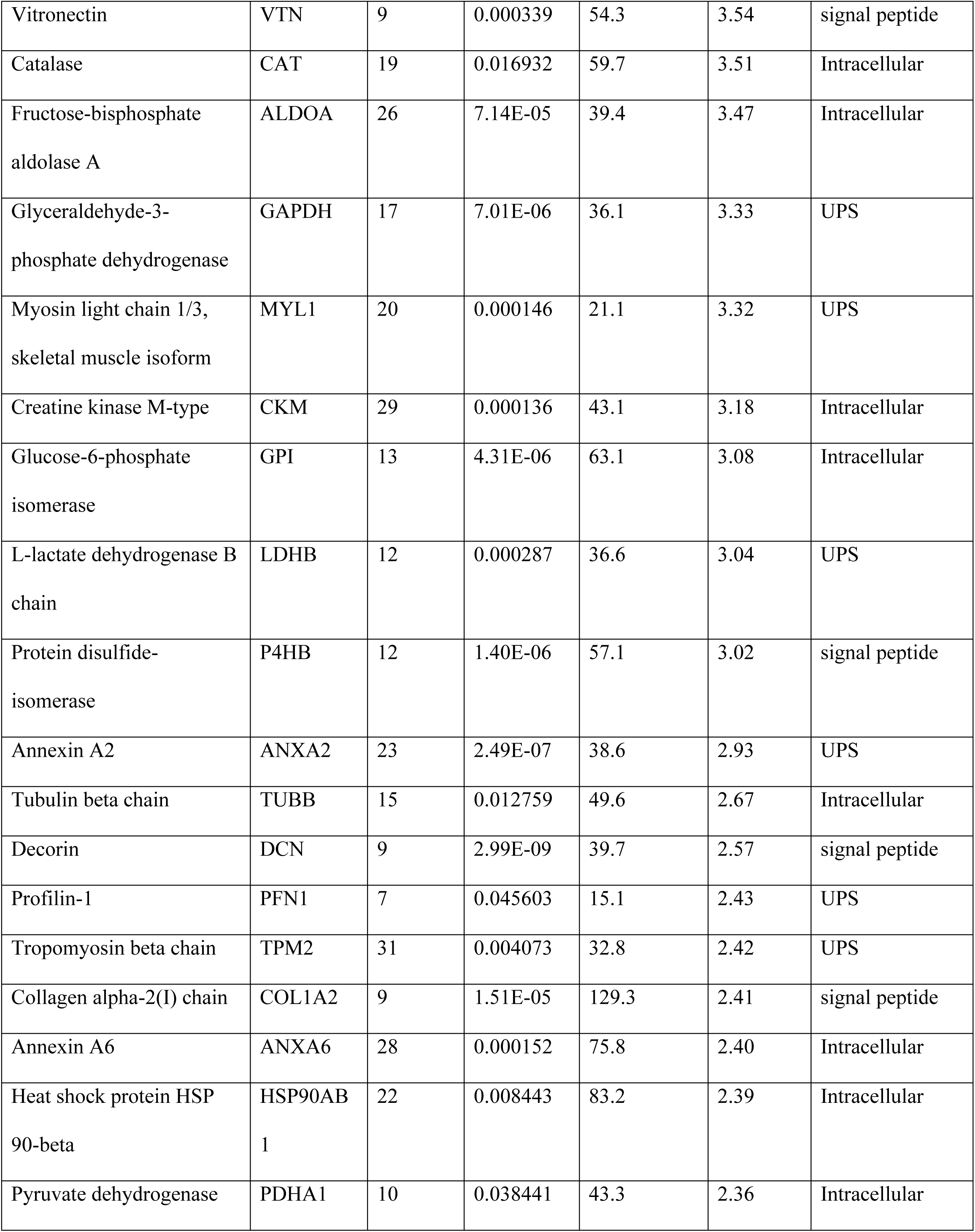
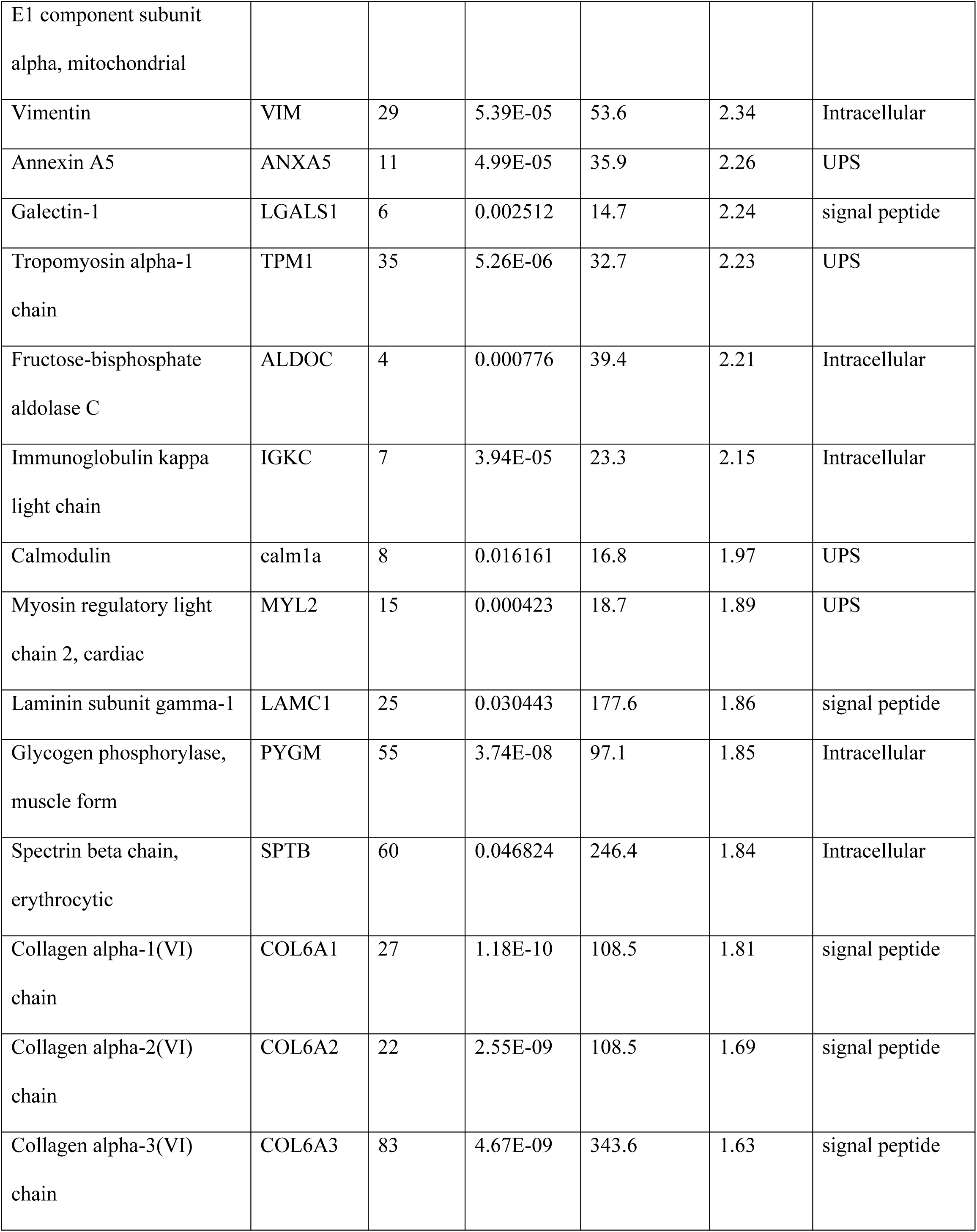
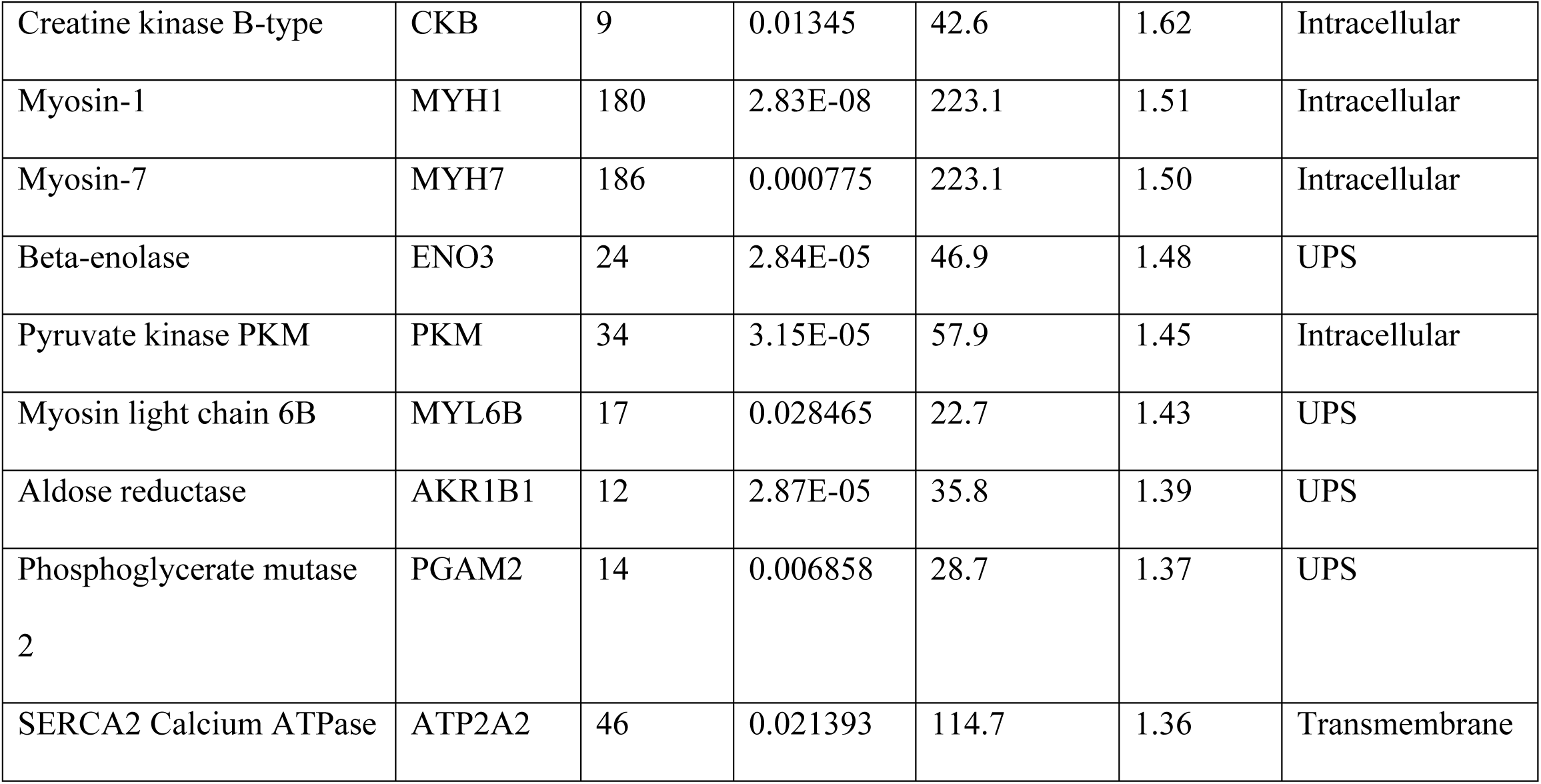
List of identified proteins that exhibit an **increased** abundance in tissue specimens from patients afflicted with **Duchenne muscular dystrophy**. Protein samples were analysed by label-free LC-MS/MS

**Table 1B.**
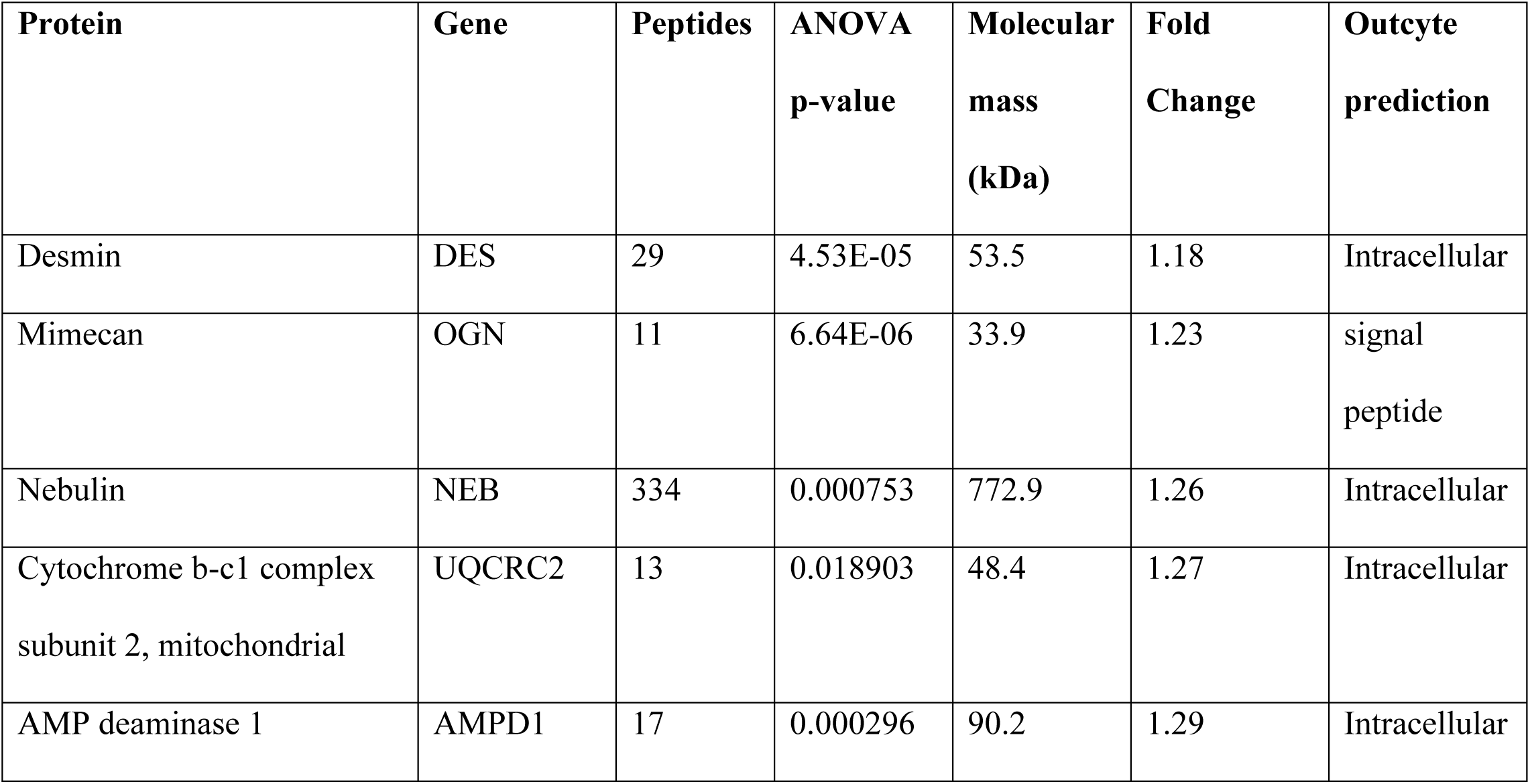
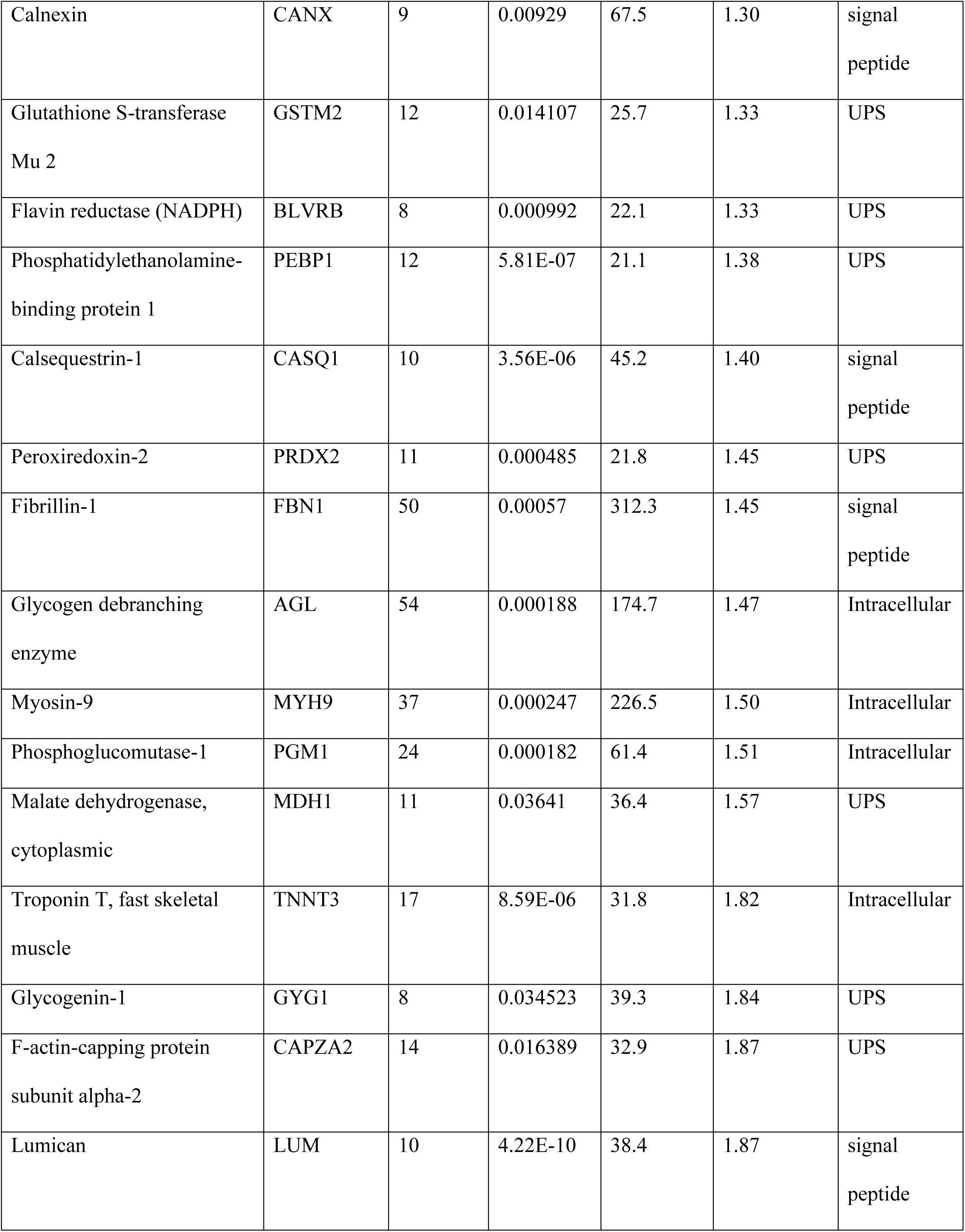
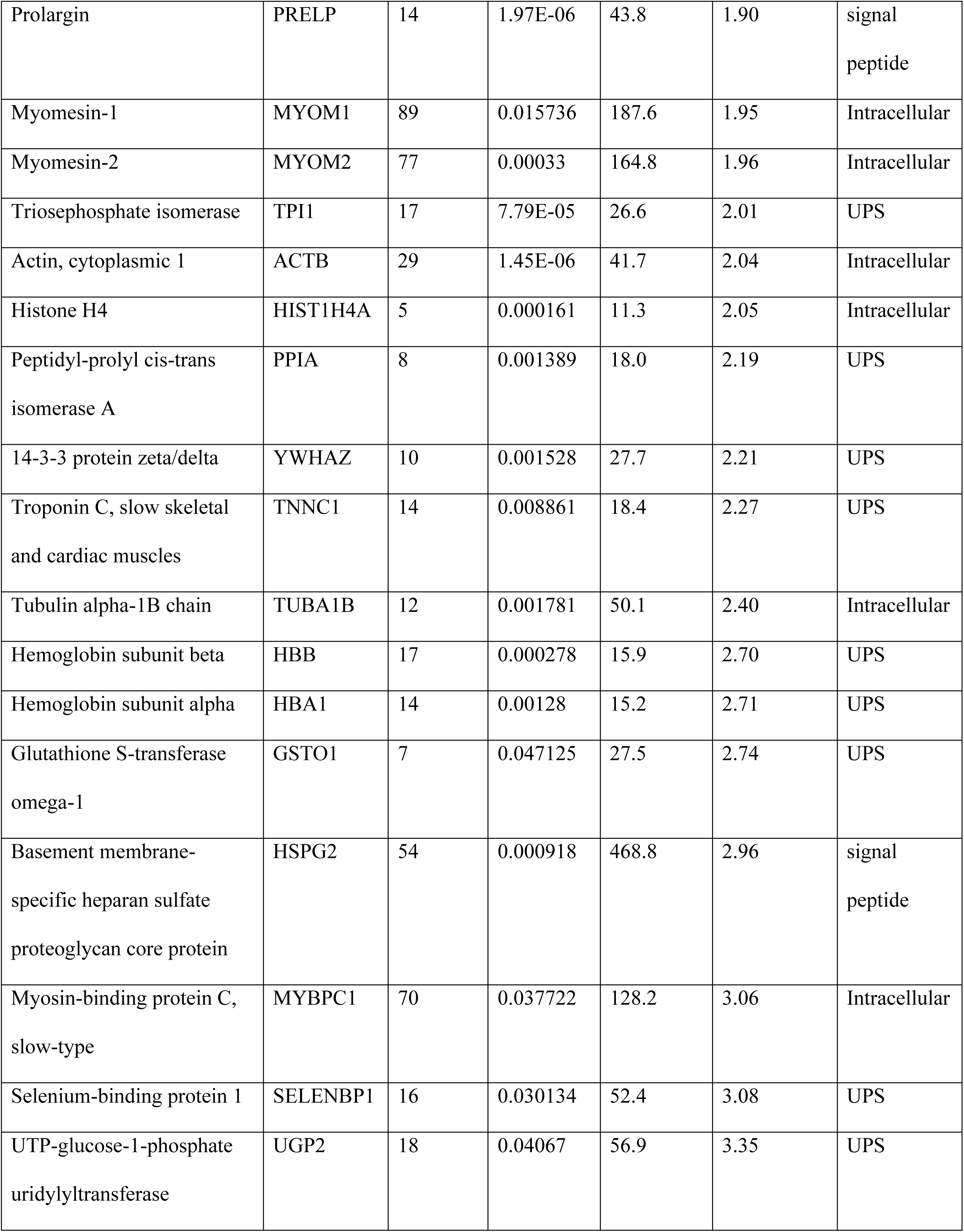
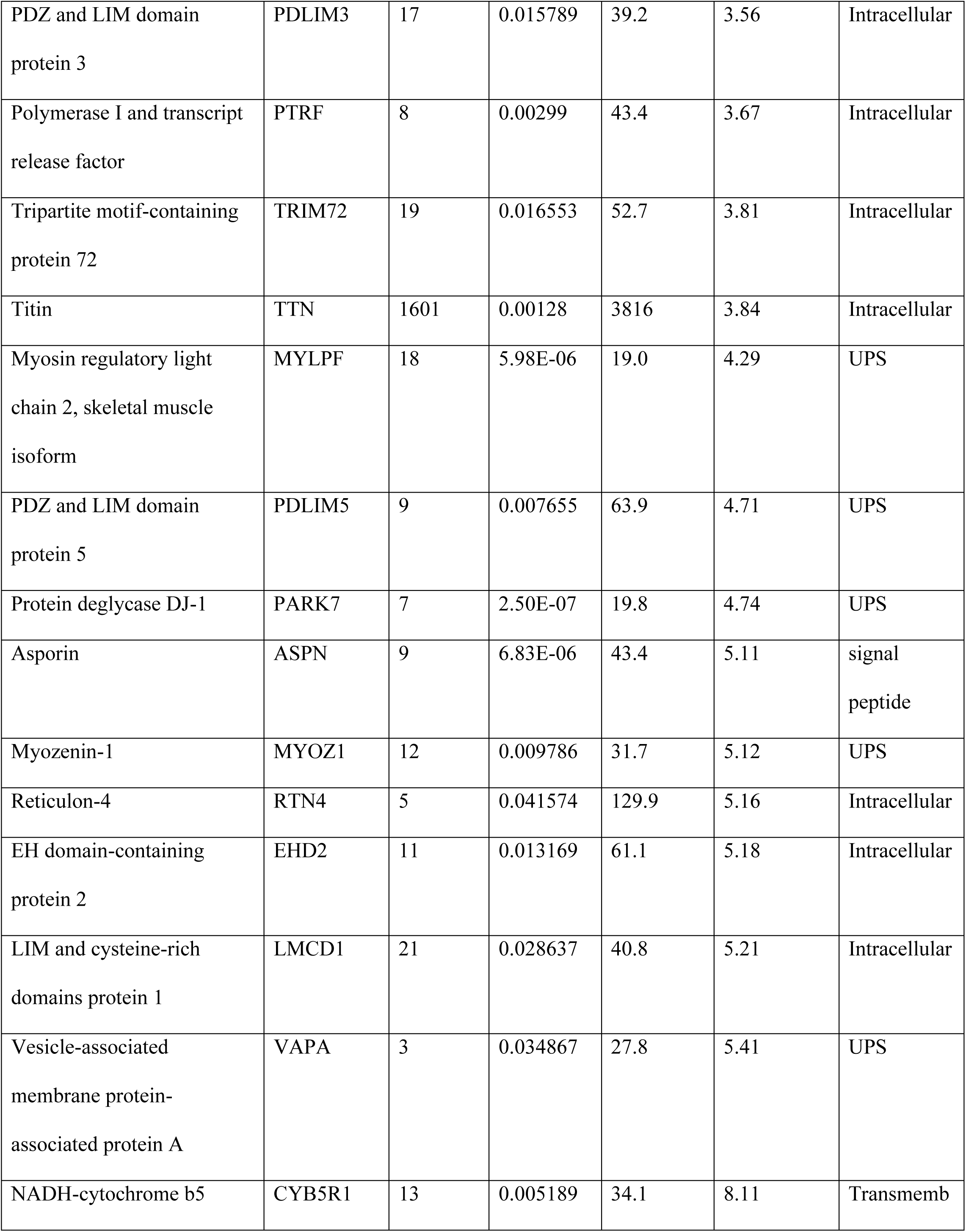
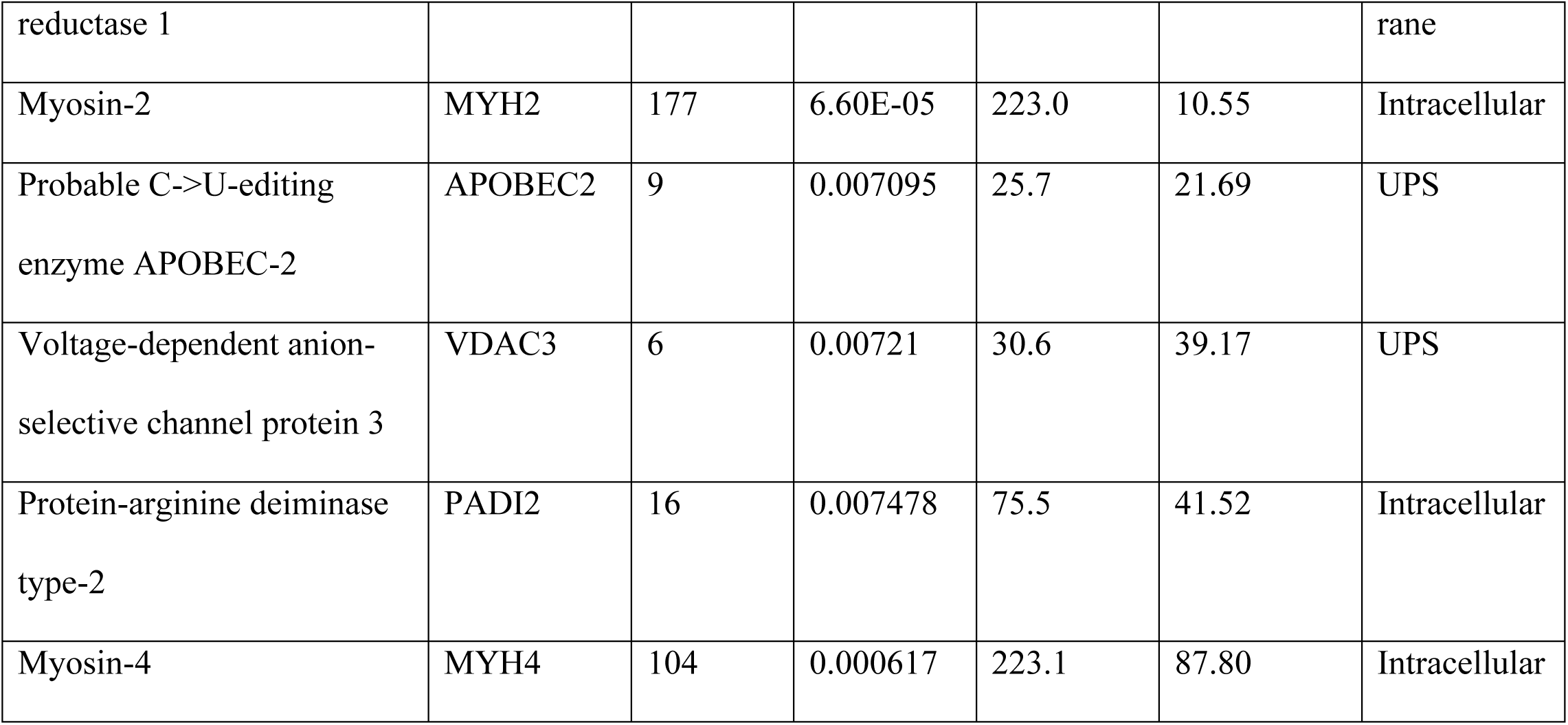
List of identified proteins that exhibit a **decreased** abundance in tissue specimens from patients afflicted with **Duchenne muscular dystrophy**. Protein samples were analysed by label-free LC-MS/MS.

**Table 2.**
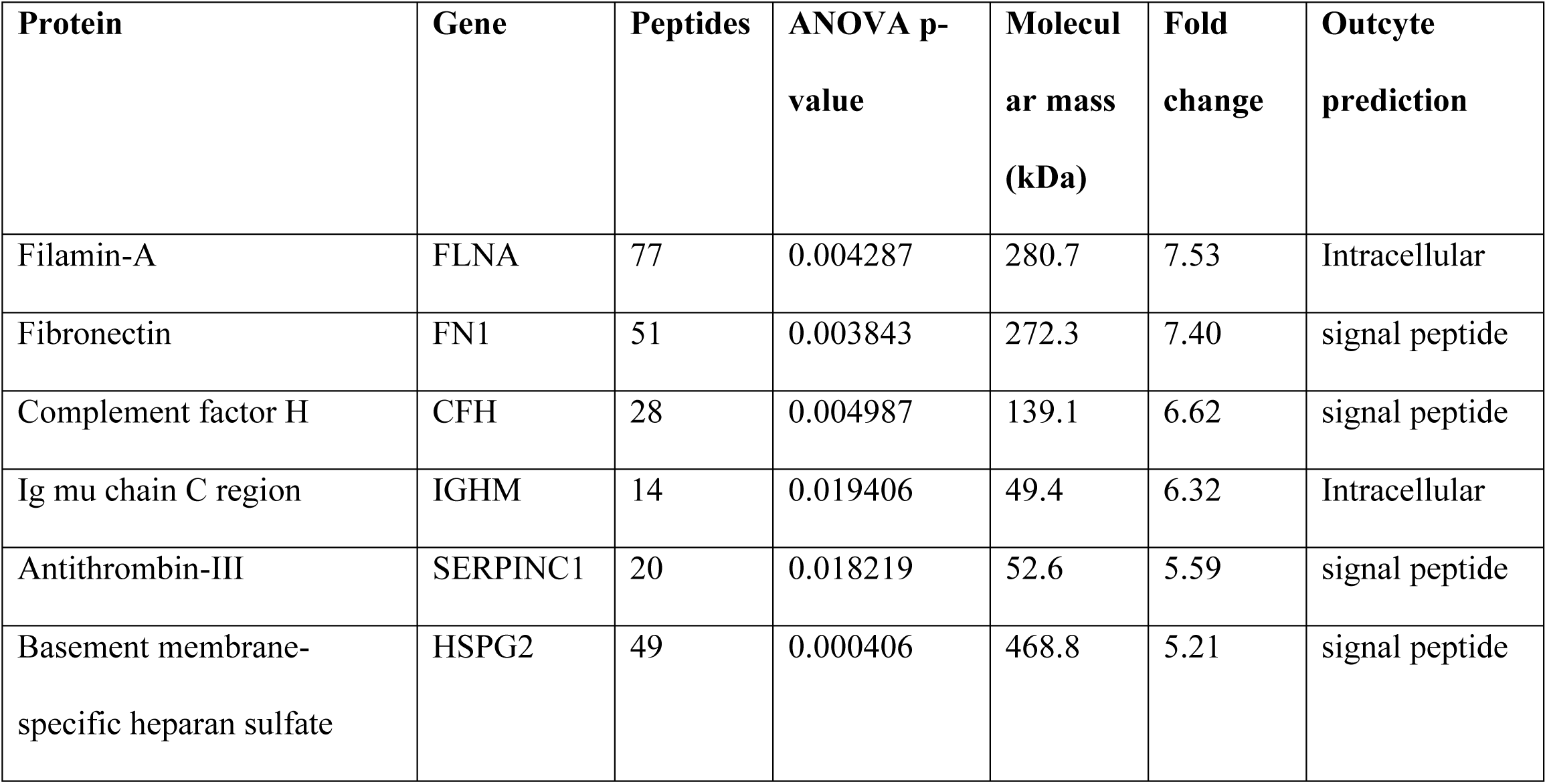
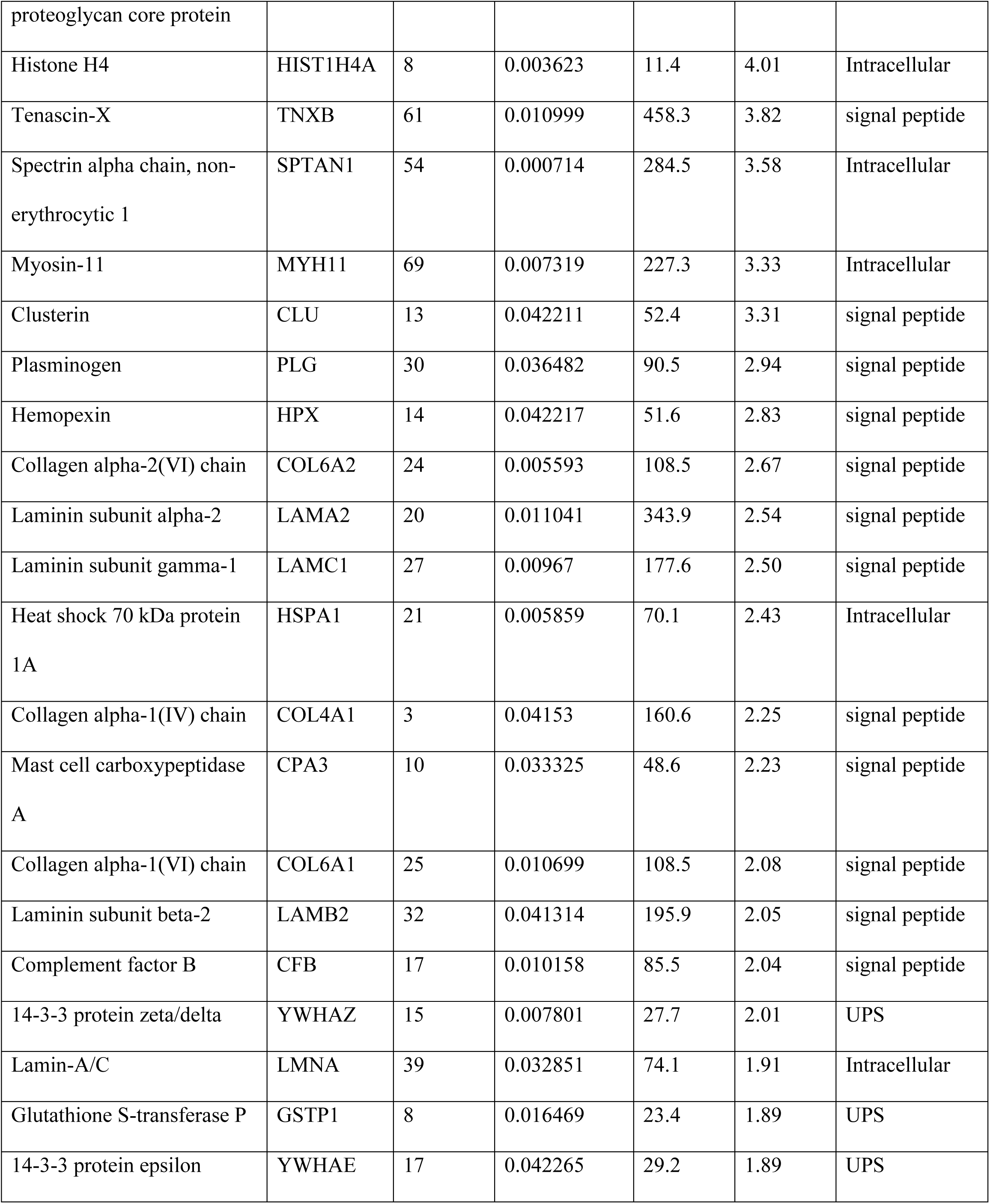
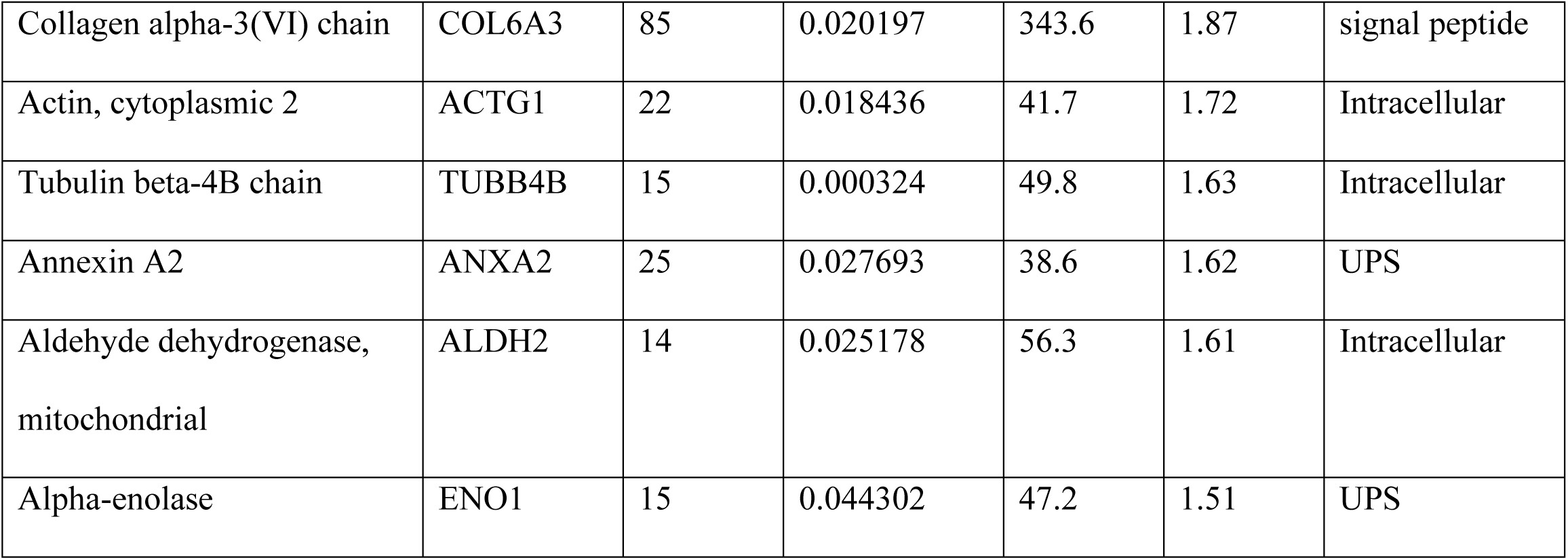
List of identified proteins that exhibit an **increased** abundance in cricopharyngeal muscle specimens from patients afflicted with **oculopharyngeal muscular dystrophy**. Protein samples were analysed by label-free LC-MS/MS.

**Table 3.** List of identified proteins that exhibit an **increased** abundance in cricopharyngeal muscle specimens from patients afflicted with **inclusion body myositis**. Protein samples were analysed by label-free LC-MS/MS.

| **Protein** | **Gene** | **Peptides** | **ANOVA p-value** | **Molecular mass (kDa)** | **Fold change** | **Outcyte prediction** |
| --- | --- | --- | --- | --- | --- | --- |
| Actin, alpha cardiac muscle 1 | ACTC1 | 31 | 0.000355 | 42.0 | 6.89 | Intracellular |
| Basement membrane-specific heparan sulfate proteoglycan core protein | HSPG2 | 68 | 0.000751 | 468.8 | 6.11 | signal peptide |
| Fibronectin | FN1 | 45 | 0.017906 | 272.3 | 5.98 | signal peptide |
| Aconitate hydratase, mitochondrial | ACO2 | 36 | 0.018057 | 85.4 | 5.26 | Intracellular |
| Citrate synthase, | CS | 12 | 0.02578 | 51.7 | 4.60 | UPS |
| mitochondrial |  |  |  |  |  |  |
| Cytochrome b-c1 complex subunit 1, mitochondrial | UQCRC1 | 19 | 0.026717 | 52.6 | 4.34 | Intracellular |
| Elongation factor Tu, mitochondrial | TUFM | 19 | 0.017793 | 49.5 | 4.31 | Intracellular |
| Histone H4 | HIST1H4A | 9 | 6.71E-05 | 11.3 | 3.75 | Intracellular |
| Plasminogen | PLG | 25 | 0.006887 | 90.5 | 3.67 | signal peptide |
| Mast cell carboxypeptidase A | CPA3 | 9 | 0.009916 | 48.6 | 3.64 | signal peptide |
| Tenascin-X | TNXB | 61 | 0.02395 | 458.3 | 3.46 | signal peptide |
| Annexin A5 | ANXA5 | 14 | 0.009982 | 35.9 | 3.43 | UPS |
| Calsequestrin-2 | CASQ2 | 11 | 0.013957 | 46.4 | 3.31 | signal peptide |
| Titin | TTN | 645 | 0.022355 | 3906.4 | 3.18 | Intracellular |
| NAD(P) transhydrogenase, mitochondrial | NNT | 24 | 0.041736 | 113.8 | 3.17 | Intracellular |
| Superoxide dismutase [Mn], mitochondrial | SOD2 | 7 | 0.026648 | 24.7 | 3.15 | UPS |
| Calreticulin | CALR | 6 | 0.011898 | 48.1 | 3.03 | signal |
|  |  |  |  |  |  | peptide |
| Annexin A1 | ANXA1 | 21 | 0.040818 | 38.7 | 2.88 | UPS |
| Collagen alpha-2(VI)<br>chain | COL6A2 | 23 | 7.24E-05 | 108.5 | 2.849 | signal<br>peptide |
| 3-ketoacyl-CoA thiolase,<br>mitochondrial | ACAA2 | 15 | 0.038653 | 41.9 | 2.83 | Intracell<br>ular |
| Plectin | PLEC | 180 | 0.013036 | 531.7 | 2.63 | Intracell<br>ular |
| Annexin A6 | ANXA6 | 30 | 0.034283 | 75.8 | 2.61 | Intracell<br>ular |
| Laminin subunit alpha-2 | LAMA2 | 48 | 0.001802 | 343.9 | 2.61 | signal<br>peptide |
| Stress-70 protein,<br>mitochondrial | HSPA9 | 19 | 0.032871 | 73.6 | 2.56 | Intracell<br>ular |
| Tubulin alpha-1B chain | TUBA1<br>B | 16 | 0.026025 | 50.1 | 2.54 | Intracell<br>ular |
| Obscurin | OBSCN | 99 | 0.036813 | 868.4 | 2.45 | Intracell<br>ular |
| Polymerase I and<br>transcript release factor | PTRF | 9 | 0.025267 | 43.4 | 2.44 | Intracell<br>ular |
| Heat shock protein HSP<br>90-beta | HSP90A<br>B1 | 25 | 0.040796 | 83.2 | 2.31 | Intracell<br>ular |
| Elongation factor 2 | EEF2 | 21 | 0.001045 | 95.3 | 2.28 | Intracell<br>ular |
| Heat shock 70 kDa | HSPA1A | 23 | 0.039811 | 70.1 | 2.28 | Intracell |
| protein 1A |  |  |  |  |  | ular |
| Desmin | DES | 33 | 0.015993 | 53.5 | 2.21 | Intracell<br>ular |
| Tubulin beta-4B chain | TUBB4<br>B | 20 | 0.000329 | 49.8 | 2.17 | Intracell<br>ular |
| Laminin subunit gamma-1 | LAMC1 | 39 | 0.013587 | 177.6 | 2.10 | signal<br>peptide |
| Glutathione S-transferase P | GSTP1 | 8 | 0.010106 | 23.3 | 2.09 | UPS |
| Tubulin alpha-4A chain | TUBA4<br>A | 16 | 0.028034 | 49.9 | 2.05 | UPS |
| Myozenin-2 | MYOZ2 | 13 | 0.012721 | 29.8 | 2.05 | Intracell<br>ular |
| Galectin-1 | LGALS1 | 6 | 0.004436 | 14.7 | 2.01 | signal<br>peptide |
| Collagen alpha-1(VI) chain | COL6A1 | 29 | 0.024197 | 108.5 | 2.01 | signal<br>peptide |
| Elongation factor 1-alpha 1 | EEF1A1 | 14 | 0.036258 | 50.1 | 1.99 | UPS |
| Myotilin | MYOT | 24 | 0.047863 | 55.3 | 1.96 | Intracell<br>ular |
| Collagen alpha-3(VI) chain | COL6A3 | 84 | 0.006678 | 343.6 | 1.96 | signal<br>peptide |
| Fibrillin-1 | FBN1 | 80 | 0.001306 | 312.3 | 1.95 | signal<br>peptide |
| Vinculin | VCL | 36 | 0.012835 | 123.8 | 1.83 | Intracellular |
| Spectrin alpha chain, non-erythrocytic 1 | SPTAN1 | 66 | 0.011121 | 284.5 | 1.70 | Intracellular |
| Myosin-9 | MYH9 | 70 | 0.02321 | 226.5 | 1.67 | Intracellular |

**Table 4A.** List of matrisome proteins **increased** in cricopharyngeal muscle from patients with **OPMD** as compared to **CTL**

| **Protein** | **Gene** | **Fold change** | **Matrisome database** |
| --- | --- | --- | --- |
| Collagen alpha-1(VI) chain | COL6A1 | 2.08 | collagens |
| Collagen alpha-2(VI) chain | COL6A2 | 2.67 | collagens |
| Collagen alpha-3(VI) chain | COL6A3 | 1.87 | collagens |
| Collagen alpha-1(IV) chain | COL4A1 | 2.25 | collagens |
| Laminin subunit alpha-2 | LAMA2 | 2.54 | glycoproteins |
| Laminin subunit beta-2 | LAMB2 | 2.05 | glycoproteins |
| Laminin subunit gamma-1 | LAMC1 | 2.50 | glycoproteins |
| Fibronectin | FN1 | 7.40 | glycoproteins |
| Tenascin-X | TNXB | 3.82 | glycoproteins |
| Basement membrane-specific heparan sulfate proteoglycan core protein | HSPG2 | 5.21 | proteoglycans |
| Annexin A2 | ANXA2 | 1.62 | ecm affiliated |
| Hemopexin | HPX | 2.83 | ecm affiliated |
| Antithrombin-III | SERPINC1 | 5.59 | regulators |
| Plasminogen | PLG | 2.94 | regulators |

**Table 4B.** List of matrisome proteins **increased** in cricopharyngeal muscle from patients with **IBM** as compared to **CTL**

| **Protein** | **Gene** | **Fold change** | **Matrisome database** |
| --- | --- | --- | --- |
| Collagen alpha-1(VI) chain | COL6A1 | 2.01 | collagens |
| Collagen alpha-2(VI) chain | COL6A2 | 2.849 | collagens |
| Collagen alpha-3(VI) chain | COL6A3 | 1.96 | collagens |
| Laminin subunit alpha-2 | LAMA2 | 2.61 | glycoproteins |
| Laminin subunit gamma-1 | LAMC1 | 2.10 | glycoproteins |
| Tenascin-X | TNXB | 3.46 | glycoproteins |
| Fibronectin | FN1 | 5.98 | glycoproteins |
| Fibrillin-1 | FBN1 | 1.95 | glycoproteins |
| Basement membrane-specific<br>heparan sulfate proteoglycan<br>core protein | HSPG2 | 6.11 | proteoglycans |
| Galectin-1 | LGALS1 | 2.01 | ecm affiliated |
| Annexin A1 | ANXA1 | 2.88 | ecm affiliated |
| Annexin A5 | ANXA5 | 3.43 | ecm affiliated |
| Annexin A6 | ANXA6 | 2.61 | ecm affiliated |
| Plasminogen | PLG | 3.67 | regulators |

**Table 4C.**
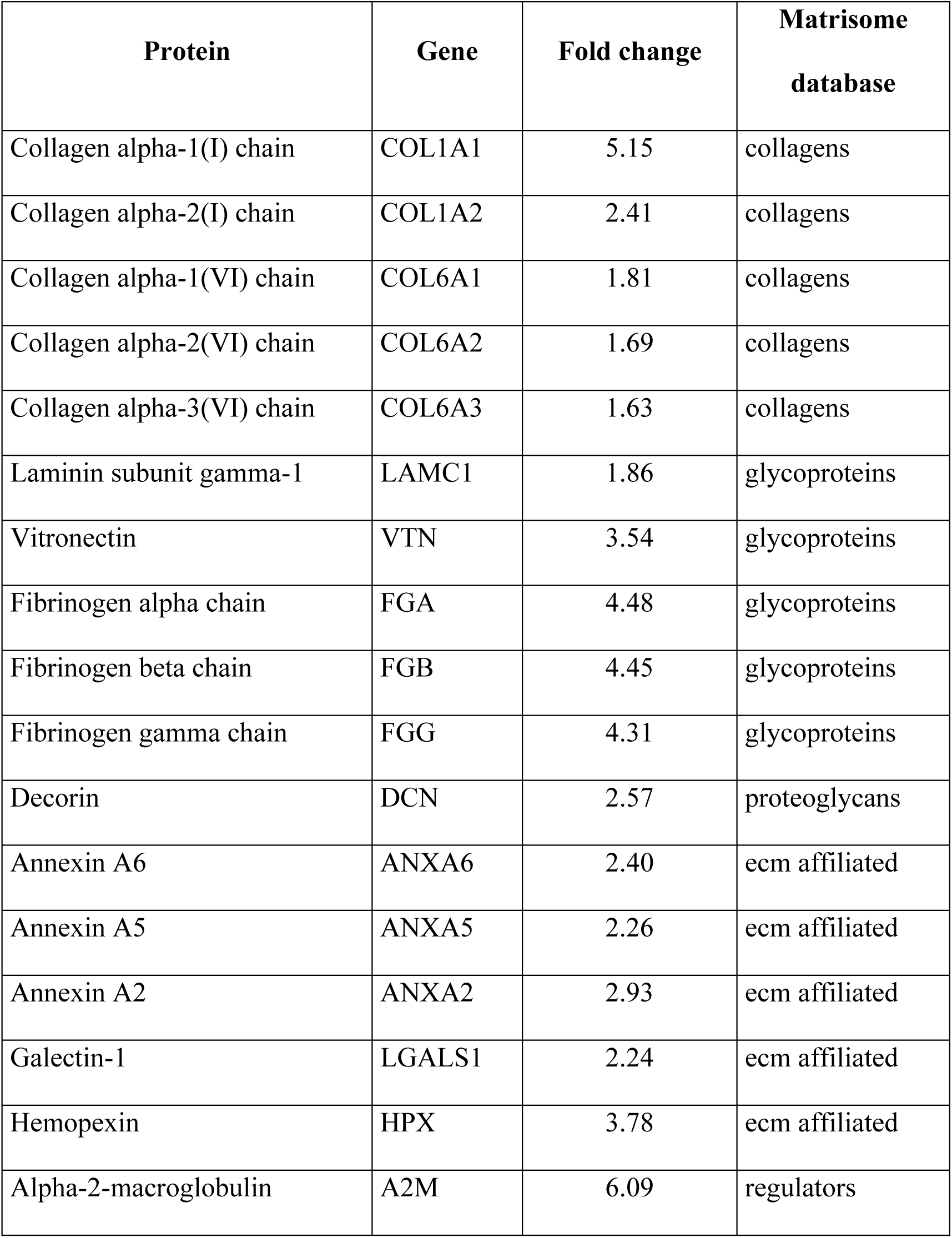
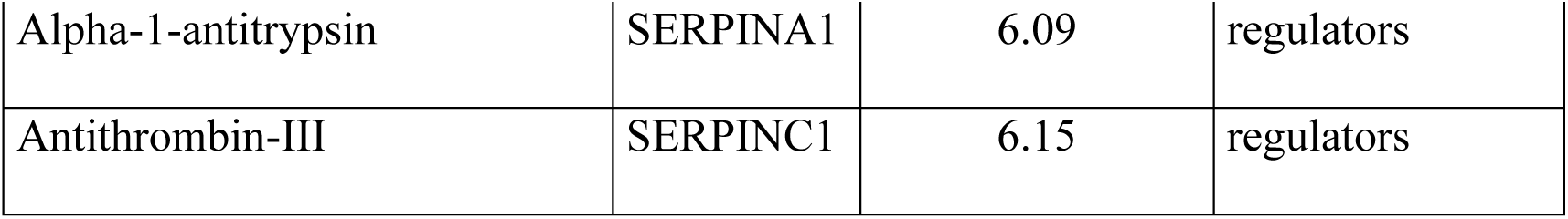
List of matrisome proteins **increased** in paraspinal muscle from patients with **DMD** as compared to **CTL**

**Table 4D.**
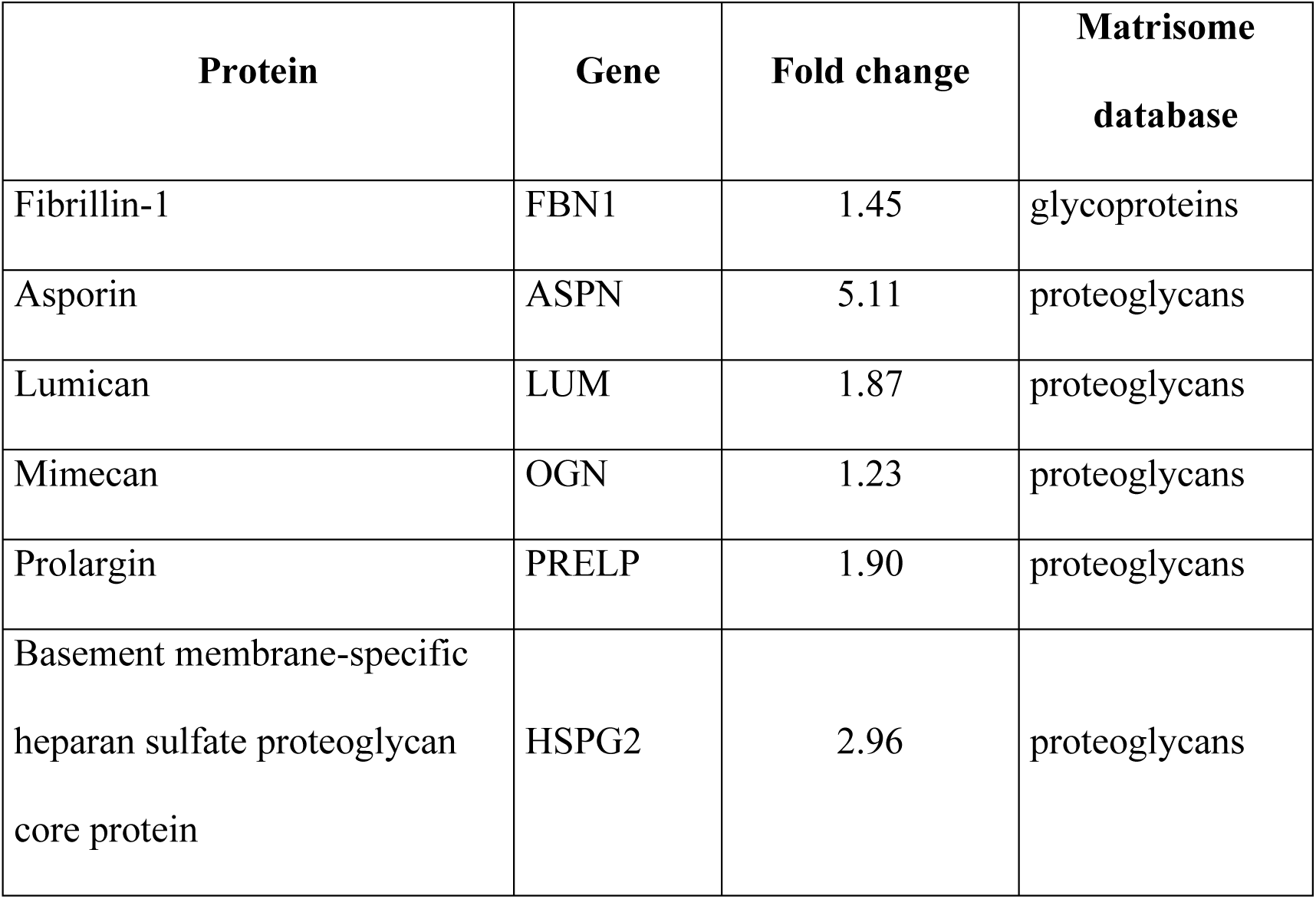
List of matrisome proteins **decreased** in paraspinal muscle from patients with **DMD** as compared to **CTL**

